# Mendelian randomisation study exploring the associations of serum folate with pan and site-specific cancers

**DOI:** 10.1101/762138

**Authors:** Kimberley Burrows, Nabila Kazmi, Philip Haycock, Konstantinos K Tsilidis, The PRACTICAL consortium, CRUK, BPC3, CAPS and PEGASUS, GECCO, CORECT and CCFR, Richard M Martin, Sarah J Lewis

**Author notes:** Members from the PRACTICAL Consortium, CRUK, BPC3, CAPS and PEGASUS are provided in the supplement. Members from the GECCO, CORECT and CCFR consortia are provided in the supplement. Corresponding author **Contact details of the corresponding authors**: Dr Kimberley Burrows, Address: Oakfield House, Oakfield Grove, University of Bristol, Bristol, BS8 2BN, UK.

## Abstract

**Background:** Epidemiological studies report evidence for an association between folate and the risk of several common cancers. However, both protective and harmful effects have been reported, and effects may differ by cancer site. Using Mendelian randomisation (MR), we investigated the causal relationships of genetically predicted serum folate with pan-cancer risk (all cancers excluding non-melanoma skin cancers); breast, prostate, ovarian, lung, and colorectal cancers; and malignant melanoma.

**Methods:** We conducted a two-sample MR analysis, using genetic instruments for serum folate to appraise the possible causal role on risk of pan-cancer and six site-specific cancers using summary statistics available from large consortia and the population-based cohort study UK Biobank (UKBB).

**Results:** There was little evidence that genetically elevated serum folate was causally associated with risk of pan-cancer or six site-specific cancers. Meta-analysis showed odds ratios (OR) per SD increase in log serum folate of 0.93 (95% CI 0.78-1.11) for breast cancer, 0.87 (95% CI 0.71-1.06) for prostate cancer, 0.84 (95% CI 0.59-1.20) for ovarian cancer, and 0.87 (95% CI 0.57-1.32) for lung cancer. The OR for colorectal cancer was 1.18 (95% CI 0.64-2.18) in large consortia analysis, while ORs for pan-cancers and malignant melanoma in UKBB were 0.88 (95% CI 0.73-1.06) and 0.56 (95% CI 0.29-1.08) respectively. The results were powered to detect modest effect sizes (>90% power (α 0.05) to detect ORs 1.2 (0.8) for the GWAS consortia) and were consistent between the two statistical approaches used (inverse variance weighted (IVW) and likelihood-based).

**Conclusions:** There is little evidence that genetically elevated serum folate may affect the risk of pan-cancer and six site-specific cancers. However, we may still be underpowered to detect clinically relevant but smaller magnitude effects. Our results provide some evidence that increasing levels of circulating folate through widespread supplementation or deregulation of fortification of foods with folic acid is unlikely to lead to moderate unintended population-wide increase in cancer risk.

**Key Messages:**

- Observational studies have identified associations between folate (both intake and circulating levels) and risk of developing site-specific cancers. However, these studies are liable to biases such as confounding, measurement error, and reverse causation.
- Using Mendelian randomisation, we appraised the causal relationships between genetically influenced serum folate levels and pan-cancer risk (all cancers excluding non-melanoma skin cancers); breast, prostate, ovarian, lung, and colorectal cancers; and malignant melanoma.
- Overall findings suggest that there is little evidence for the causal associations between genetically influenced serum folate and risk of pan-cancer and six site-specific cancers.
- We provide some evidence that increasing levels of circulating folate through widespread supplementation or deregulation of fortification of foods with folic acid is unlikely to lead to moderate unintended population-wide increase in cancer risk.

## Introduction

Folate is an essential B vitamin found in foods such as dark leafy green vegetables, liver and legumes. Serum folate reflects recent folate intake and is the earliest biomarker to detect folate status[1]. Folic acid, the synthetic form of folate, is available as a dietary supplement and is used to fortify food such as bread flour in over 80 countries worldwide[2].

Folate has an essential role in the synthesis and methylation of DNA and is a crucial co-factor in one-carbon metabolism together with other B vitamins such as vitamins B2, B6, and B12[3]. In developing foetuses, insufficient folate increases the risk of neural tube defects, including spina bifida and anencephaly[4,5]. In adults, insufficient folate can lead to anaemia[6]. Low folate levels may contribute to carcinogenesis through aberrations in DNA methylation and uracil misincorporation, leading to DNA instability[7]. However, folic acid supplementation is shown to have tumour-promoting effects in mouse models[8].

Epidemiological studies exploring associations of folate with the risk of developing site-specific cancers have been inconsistent. For instance, total folate, dietary folate and serum folate levels have been reported to have no associations with breast cancer[9,10], whilst in contrast, a meta-analysis of 26 case-control studies reports protective effects of higher dietary folate intake[11]. Likewise, some meta-analyses suggest positive associations between serum folate and prostate cancer[12], while others suggest little evidence of associations with folate intake[13,14]. These inconsistencies are also present for studies examining folate and colorectal cancer[15,16]. Much of the observational studies to date are limited due to small study sample sizes, measurement error, heterogeneity of the exposure measurement (dietary intake vs. supplement intake vs. circulating levels), timing of folate measurement (leading to possible reverse causation), and the use of data from both pre-and post-folic acid fortification study populations[17].

Several randomised control trials (RCTs) have been conducted exploring the effects of folic acid supplementation on a range of primary outcomes while having also recorded incident cancers. A 2013 pooled analysis of folic acid supplementation recorded 3713 cancer incidents in around 50 000 participants with a weighted average treatment period of 5.2 years (range 1.8 to 7.4 years). The meta-analysis reported little evidence that folic acid treatment increased (or decreased) risk of overall cancer or cancers of the colorectum, lung, ovaries, breast, malignant melanoma, or prostate compared to placebo[18]. However, these trials are limited by the small number of incident cancer cases and the short duration of treatment time during the trials.

Mendelian randomisation (MR) is an instrumental variable analytical approach which utilises common genetic variants as instruments to proxy potentially modifiable risk factors. The aim of MR is to elucidate the causal effects of these risk factors on disease outcomes of interest[19,20]. Germline genetic variants are randomised and fixed at conception, enabling MR analysis to mitigate the major biases of observational studies such as residual confounding, measurement error and reverse causation. The aim of the current study was to apply MR within a two-sample framework to elucidate the causal associations of serum folate with pan-cancer risk; cancers of the breast, prostate, ovaries, lung, and colorectum; and malignant melanoma.

## Materials and Methods

### Genetic instrument selection for serum folate

We conducted a search of published genome-wide association studies (GWAS) using MR-Base[21] and PubMed (https://www.ncbi.nlm.nih.gov/pubmed/). Studies with single nucleotide polymorphisms (SNPs) that were robustly associated at P-value <5×10^−8^ with serum folate levels and involving participants of European ancestry were prioritised. We identified a moderately sized GWAS of serum folate in a healthy, young adult Irish population consisting of 2232 individuals with full summary statistics available[22]. Five SNPs (rs1801133, rs1999594, rs12085006, rs7545014, and rs7554327) from Shane *et al*.[22] located within a 100kb region around the methylenetetrahydrofolate reductase (*MTHFR*) gene were identified as potential instruments. We excluded rs12085006 and rs7554327 as they are in near-perfect linkage disequilibrium (LD) with rs1999594 (R^2^ 1.00) and rs7545014 (R^2^ 0.99) respectively. Detailed information on the selected genetic instruments is provided in Table 1.

**Table 1.** Genetic variants included in the instrumental variable and their associations with serum folate.

| Variant | chr:pos | Locus | Alleles |  | EAF | Per-allele estimate |  |  | R<sup>2</sup> |
| --- | --- | --- | --- | --- | --- | --- | --- | --- | --- |
|  |  |  | Effect | Other |  | Beta<sup>++</sup> | SE<sup>+</sup> | P-value |  |
| rs1801133 | 1:11778965 | *MTHFR* | C | T | 0.66 | 0.062 | 0.009 | 2.82E-11 | 0.020 |
| rs7545014 | 1:11857240 | *LOC390997* | C | T | 0.56 | 0.050 | 0.009 | 1.01E-08 | 0.015 |
| rs1999594 | 1:11881803 | *RNU5E* | T | C | 0.45 | 0.050 | 0.009 | 1.43E-08 | 0.014 |

chr, chromosome; pos, position (Build 38); Locus, nearest gene reported [Shane et al. 2017]; EAF, effect allele frequency where the effect allele is the folate increasing allele; SE, standard error; MTHFR, Methylene tetrahydrofolate Reductase; LOC390997, SET Binding Factor 1 Pseudogene; RNU5E, RNA-U5E Small Nuclear 1; R<sup>2</sup>, proportion of variance explained; \* Linear regression adjusted for age and sex; + Regression coefficients were re-scaled to represent SD change per each additional effect allele.

### Data on the genetic epidemiology of cancers

We retrieved summary statistics of the genetic effects for the selected instruments on the risk of site-specific cancers from large, recently published GWAS. Four large consortia had publicly available summary statistics for breast cancer (BCAC - Breast Cancer Association Consortium), prostate cancer (PRACTICAL - Prostate Cancer Association Group to Investigate Cancer Associated Alterations in the Genome), ovarian cancer (OCAC - Ovarian Cancer Association Consortium), and lung cancer (ILCCO - International Lung Cancer Consortium). Summary statistics were made available for colorectal cancer from the Genetic and Epidemiology of Colorectal Cancer Consortium (GECCO), the Colorectal Cancer Transdisciplinary Study (CORECT), and the Colon Cancer Family Registry (CCFR) consortia (GECCO-CORECT-CCFR). In Supplementary methods we further describe each of these datasets. Information on quality control, imputation and statistical analysis for each GWAS has been previously reported[23–27].

The UK Biobank is a population-based cohort study consisting of approximately 500 000 middle-aged participants, who were recruited between 2006 and 2010 from across the UK[28,29]. We performed GWAS for cancers of the breast, prostate, ovaries, lung, colorectum and malignant melanoma identified via linkage to the UK Cancer Registry. Cases were defined as having a cancer diagnosis occurring either before or after enrolment to the UKBB study. A list of the ICD09 and ICD10 codes used to define each site-specific cancer are included in Supplementary Table S1. We also performed GWAS for pan-cancer in UKBB using data for every cancer site reported (excluding non-melanoma skin cancers). Further details on the definition of cases and controls, GWAS and statistical analysis can be found in the Supplementary Methods and Supplementary Table S1. All three instruments for serum folate were available in each of the GWAS consortia and in UKBB.

### Mendelian randomisation analysis

We conducted two-sample MR analyses to appraise the causal relationships between serum folate and the risk of pan-cancer and six site-specific cancers (breast, prostate, ovarian, colorectal, lung and malignant melanoma)[30].

We re-scaled the SNP-folate effect estimates to the standard deviation (SD) scale to represent an SD change in log_10_ transformed serum folate with each additional effect allele (see supplementary methods). We harmonised the SNPs so that the effect alleles were the serum folate increasing alleles.

The three SNPs are located within a 100 kb region around the *MTHFR* gene on chromosome 1 and are in weak LD with each other (all R^2^ <0.45) (see Supplementary Table S2). The use of multiple correlated SNPs introduces bias of over precision of the overall causal effect estimates. To mitigate this, we used extensions of the fixed-effect inverse variance weighted (IVW) method and the likelihood-based approach to account for the correlation structure between the SNPs[31,32].

Fixed-effects IVW meta-analysis was performed to pool the MR estimates from the GWAS consortia studies and UKBB for the following cancers: breast, prostate, ovarian, and lung. The GECCO-CORECT-CCFR consortia GWAS of colorectal cancer included samples from the UKBB and so it was not appropriate to meta-analyse these results. Cochran’s Q statistic was used to assess heterogeneity between studies.

### Sensitivity analyses

The validity of the effect estimates and interpretation in MR analyses are reliant on the following assumptions[34]: i) the selected genetic instruments are robustly associated with serum folate; ii) the genetic instruments affect cancer only through their effect on serum folate; and iii) the instruments are independent of any confounders of the association between serum folate and cancer.

To evaluate the first MR assumption, we estimated the variance in serum folate explained (R^2^) by each SNP as well as the strength of the instruments represented by the F-statistic. The R^2^ and the F-statistic can be used to evaluate the strength of our instruments and to indicate weak instrument bias[35]. Derivation of the R^2^ and the F-statistic is given in the Supplementary methods. To evaluate potential violation of the second and third assumption, we performed look-ups for each of our instruments using the MR-Base PheWAS (http://phewas.mrbase.org/) tool to determine the presence of associations with secondary phenotypes that could be potential confounders of the association. Due to the limited number of folate SNPs, and their correlation, we were unable to assess potential violations of the second assumption of MR (no horizontal pleiotropy) using statistical methods (MR-Egger, weighted median and mode estimators).

Cochran’s Q statistic was calculated to assess heterogeneity across SNPs in the causal estimate[36]. Where there was evidence of heterogeneity (P-value <0.05), a (multiplicative) random-effects IVW and maximum likelihood MR analysis[37] was performed.

To further elucidate the potential impact of using correlated SNPs as an instrument we derived causal estimates for each individual SNP by calculating the ratio of coefficients (Wald ratios)[38]. The corresponding SEs were derived using the delta method[39]. In addition, we explored systematically whether an individual SNP was driving the main MR association results by performing a leave-one-out analysis, whereby IVW estimates are derived iteratively by excluding each SNP in turn.

### Statistical power

Power calculations were performed using the online tool mRnd (http://cnsgenomics.com/shiny/mRnd/) as described previously[40]. We had >90% power to detect modest effect sizes (OR 1.2 or its inverse 0.8) in our MR analysis for consortia studies of breast (BCAC), prostate (PRACTICAL), ovarian (OCAC), lung (ILCCO), and colorectal (GECCO-CORECT-CCFR) cancers as well as pan-cancers from UKBB. For UKBB, where cases were not enriched within the dataset, power to detect an OR of 1.2 ranged from 34% to >99%. Sample sizes for each cancer and detailed power calculations for a range of effect sizes are shown in Supplementary Table S3.

Analyses were conducted in R software version 3.5.1 using TwoSampleMR and MRInstruments[21], MendelianRandomization[41], meta, and matafor R packages. All reported P-values are two-tailed.

## Results

Table 1 describes the associations for each SNP comprising our instrument with serum folate, after rescaling to the SD scale. In total, the genetic instruments explained 4.9% of the variance in serum folate levels. The corresponding F-statistic (113.8) suggests that weak instrument bias was unlikely[35].

### Mendelian randomisation estimates for the association between serum folate and cancer

Pan-cancer and cancers of the breast, prostate, ovaries, lung and malignant melanoma showed consistent inverse effects using both the IVW method and the likelihood-based approach. In contrast, the colorectal cancer causal estimates suggest increasing risk with increasing serum folate. These results were concordant between those of the consortia studies and UKBB. However, our estimates were imprecise with 95% confidence intervals crossing the null suggesting that there is little evidence of causal associations (Table 2 and Figure 1A). As effect estimates were very similar between the IVW method and the likelihood-based approach, all subsequent analyses utilise the IVW estimates.

**Table 2.** Mendelian randomisation estimates between genetically elevated serum folate and risk of cancer.

| Cancer type | Study | Inverse variance weighted estimate |  |  |  | Maximum likelihood estimate |  |  |  | Q | Phet |
| --- | --- | --- | --- | --- | --- | --- | --- | --- | --- | --- | --- |
|  |  | OR | LCI | UCI | P value | OR | LCI | UCI | P value |  |  |
| **Pan-cancers** | UKBB | 0.880 | 0.729 | 1.062 | 0.183 | 0.876 | 0.722 | 1.064 | 0.182 | 2.053 | 0.358 |
| **Breast** | BCAC | 0.939 | 0.784 | 1.124 | 0.489 | 0.936 | 0.778 | 1.125 | 0.481 | 2.876 | 0.238 |
|  | UKBB* | 0.825 | 0.412 | 1.653 | 0.587 | 0.803 | 0.382 | 1.688 | 0.563 | 7.991 | 0.018 |
| **Prostate** | PRACTICAL** | 0.860 | 0.681 | 1.085 | 0.203 | 0.850 | 0.666 | 1.086 | 0.194 | 4.195 | 0.123 |
|  | UKBB | 0.889 | 0.581 | 1.360 | 0.587 | 0.886 | 0.574 | 1.365 | 0.582 | 1.845 | 0.398 |
| **Ovarian** | OCAC | 0.862 | 0.593 | 1.255 | 0.439 | 0.860 | 0.587 | 1.258 | 0.437 | 1.350 | 0.509 |
|  | UKBB | 0.672 | 0.215 | 2.103 | 0.494 | 0.657 | 0.202 | 2.132 | 0.484 | 3.478 | 0.176 |
| **Lung** | ILCCO | 0.810 | 0.488 | 1.343 | 0.414 | 0.809 | 0.486 | 1.347 | 0.415 | 0.377 | 0.828 |
|  | UKBB | 1.018 | 0.471 | 2.201 | 0.963 | 1.019 | 0.464 | 2.239 | 0.962 | 2.608 | 0.272 |
| **Colorectal** | GECCO*** | 1.178 | 0.635 | 2.184 | 0.604 | 1.224 | 0.614 | 2.440 | 0.892 | 12.594 | 0.002 |
|  | UKBB | 1.449 | 0.851 | 2.465 | 0.172 | 1.455 | 0.846 | 2.505 | 0.176 | 0.851 | 0.654 |
| **Malignant melanoma** | UKBB | 0.562 | 0.293 | 1.079 | 0.083 | 0.561 | 0.287 | 1.094 | 0.090 | 0.346 | 0.841 |

OR, odds ratio; LCI, lower 95% confidence interval; UCI, upper 95% confidence interval; Cochran's Q and P<sub>het</sub> is the P-value for heterogeneity between instrumented SNP causal estimates in the IVW analysis; OR and 95% CI reflect the causal risk estimate of cancer per genetically predicted standard deviation increase in log<sub>10</sub> serum folate; \* causal estimators are derived using random effects for both IVW and maximum likelihood models; + the PRACTICAL Consortium, CRUK, BPC3, CAPS and PEGASUS and the GECCO, CORECT and CCFR consortia.

**Figure 1.**
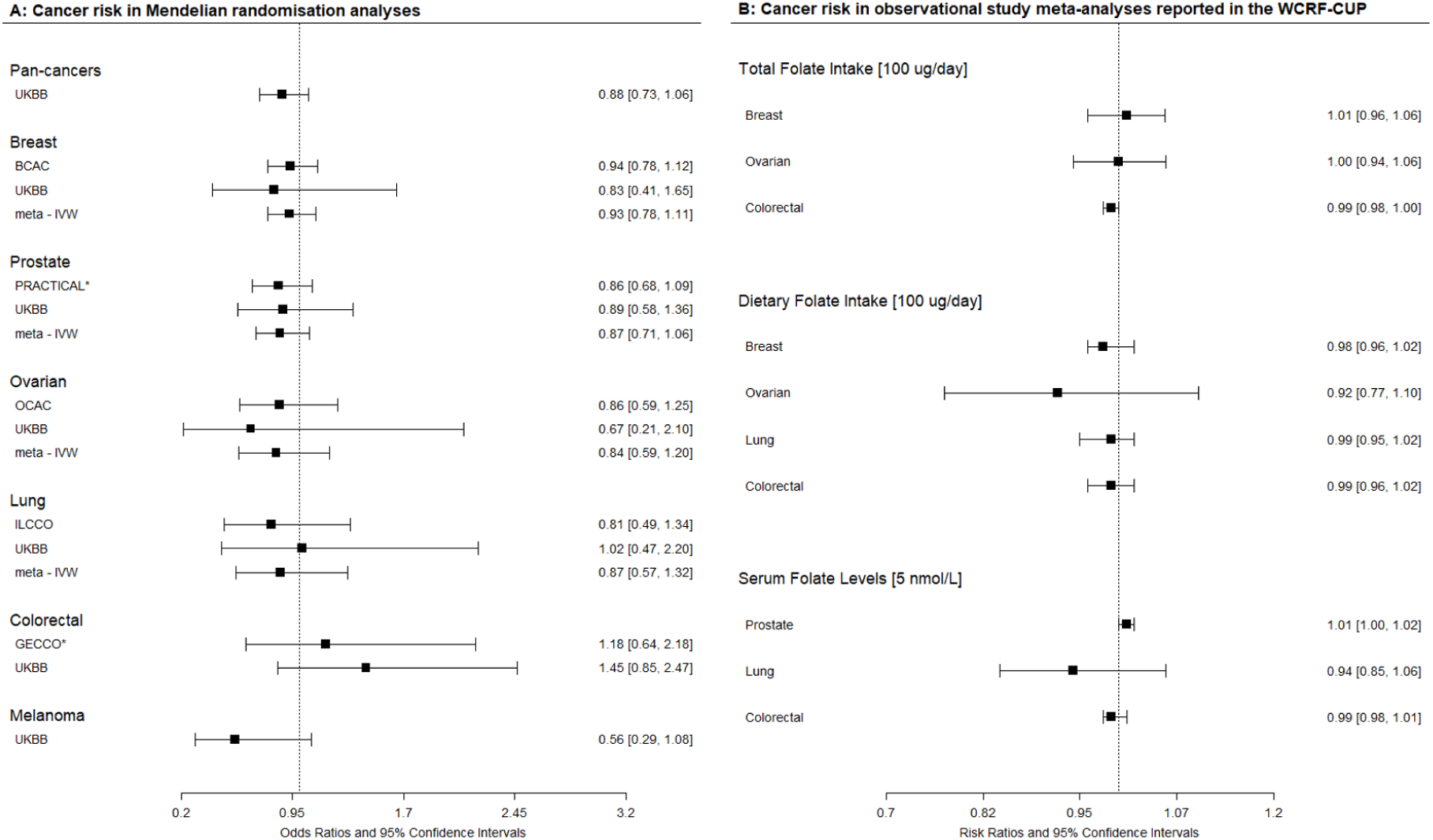
Forest Plot of Mendelian randomisation causal association estimates between serum folate and cancers. The odds ratios (OR) were derived using the inverse variance weighted method and correspond to a 1 SD increase in log10 serum folate levels. Meta – IVW correspond to the fixed effects IVW meta-analysis results; Observational studies report risk ratios; the details of which are in Supplementary Table S7. * the PRACTICAL Consortium, CRUK, BPC3, CAPS and PEGASUS and the GECCO, CORECT and CCFR consortia.

Figure 1A shows a forest plot depicting our MR causal estimates for each cancer study as well as the combined effects using fixed-effects IVW meta-analysis for breast, prostate, ovarian, and lung cancer. Combined estimates were concordant in magnitude and direction of effect to those of the individual studies. Again, there was little evidence of causal associations with confidence intervals crossing the null. In addition, there was little evidence of heterogeneity between the studies in each meta-analysis (all Cochran’s Q P-values >0.6; Supplementary Table S5).

### Sensitivity analyses and evaluation of Mendelian randomisation assumptions

Overall, there was little evidence of heterogeneity of effect estimates between each of the three serum folate SNPs (Cochran’s Q P-value >0.1) in our MR analyses. This is with the exception of breast cancer in UKBB (Cochran’s Q P-value 0.02) and colorectal cancer in GECCO-CORECT-CCFR (Cochran’s Q P-value 0.002). Random effects IVW and likelihood approach are therefore reported for these two cancers (Table 2). Supplementary Figure S1 shows scatter plots of associations between serum folate SNPs and the risk of each of the cancer studies analysed. Results for individual single SNP MR analysis (using Wald ratios) are provided in Supplementary Table S4. There is no strong evidence for causal associations between any of the individual SNPs and cancer. Overall, the effect estimates were concordant with that of the IVW MR analyses.

Leave-one-out analysis displayed concordant direction of effect estimates for all cancer outcomes except colorectal cancer, suggesting that it is unlikely that any individual SNP was driving the IVW MR results (Supplementary Figure S2). For colorectal cancer, SNP rs1999594 showed a negative effect estimate for cancer risk compared to the positive effect estimates for the other two SNPs. This may explain some of the heterogeneity between the SNPs, the wider IVW MR confidence intervals and the discordant Wald ratios between the SNPs.

After lookup within the MR-Base PheWAS database, we found some evidence from GWAS that the three SNPs were associated with additional phenotypes at genome-wide significance (P-value <1×10^−5^) (Supplementary Table S6). All three SNPs were associated with blood cell traits; rs1801133 is associated with mean corpuscular haemoglobin, mean corpuscular volume, plateletcrit (a measure of total platelet mass), and red cell distribution width[42]. Whilst rs7545014 and rs1999594 are associated with plateletcrit and platelet count[42]. In addition, rs1801133 is associated with several vascular phenotypes including diastolic blood pressure and hypertension in UKBB as well as birthweight of first child and hip circumference. Rs1999594 is associated with diastolic blood pressure and rs7545014 is associated with the operative procedure to excise umbilicus; both within UKBB.

We compared the direction of our meta-analysed MR causal estimates to those of meta-analyses of observational studies reported in the World Cancer Research Fund Continuous Update Project (WCRF-CUP)[43]. The WCRF-CUP aimed to systematically review and meta-analyse observational studies and RCTs associating nutritional risk factors with site-specific cancers. Figure 1B illustrates a forest plot of effect estimates (RR) and 95% confidence intervals of observational effect estimates for serum folate, dietary folate intake and total folate intake (diet and supplements) where data is available. While we cannot directly compare the magnitude of effect owing to the differences in measures and units used, we can observe whether there are comparable protective or adverse effects on the site-specific cancers. In total, the meta-analyses of serum folate, dietary folate intake and total folate intake showed little evidence of associations with site-specific cancers. Details of the observational study results are given in Supplementary Table S7.

## Discussion

We found no strong evidence that genetically elevated serum folate was causally associated with pan-cancer and six site-specific cancers. Although we found little evidence of causal associations, the MR effect estimates tended towards those of being protective for cancer risk with increasing serum folate levels.

### Breast cancer

In line with our findings for breast cancer, the latest WCRF-CUP reported little evidence of association between dietary and total folate intake with breast cancer risk[43] (Supplementary Table S7). Subsequent meta-analyses have also reported concordant results to those reported in our MR study. The European Prospective Investigation into Cancer and Nutrition (EPIC) cohort recently reported protective effects of similar magnitude to our MR analysis of plasma folate on the risk of breast cancer albeit with little statistical evidence (OR 0.93; 95% CI 0.83-1.05)[44].

A recent systematic review and meta-analysis have demonstrated a U-shaped dose-effect relationship between dietary folate intake and breast cancer risk in prospective studies. Daily intake of folate between 153 and 400 µg showed a reduced breast cancer risk compared to those with low folate intake (<153 µg), but not for those >400 µg[45]. We were unable to explore potential non-linear relationships within our MR study owing to lack of individual-level data.

More recently, authors included within our study published findings for an MR of circulating concentrations of micro-nutrients and risk of breast cancer in BCAC[46]. In common with our study, SNP rs1801133 was included within the serum folate instrument. Per 1 SD increase in serum folate (nmol/L), the authors reported an OR 1.06 (95% CI 0.94-1.20). The effect estimates are at odds with those we have presented (OR 0.94; 95% CI 0.78-1.12). However, both are imprecise and have confidence intervals that overlap. When comparing the causal effect estimates of rs1801133 alone; we see a more comparative causal estimate (OR 1.03; 95% CI 0.89-1.18 Papadimitriou *et al*.[46] vs. OR 1.04; 95% CI 0.82-1.26). It is important to note that our MR study reports causal effect estimates per 1 SD increase in log_10_ serum folate (nmol/L) while the estimates for Papadimitriou *et al* are reported per 1 SD increase in serum folate on the natural scale (nmol/L).

### Prostate cancer

In a pooled nested case-control study (6875 cases, average follow-up of 8.9 years) higher (vs. lowest fifth) serum folate was strongly associated with increased risk of prostate cancer (OR 1.13; 95% CI 1.02-1.26)[12]. This positive relationship was further demonstrated in another meta-analysis (OR 1.43; 95%CI 1.06-1.93)[13]. In the WCRF-CUP[43] meta-analysis there was little evidence to support such associations with serum folate (5938 cases, RR 1.01 per 5 nmol/L; 95% CI 1.00-1.02); in line with our MR study. These inconsistencies may arise due to study design; nested case-control studies with relatively short follow up times, as well as appreciably fewer cases compared to our MR study.

In a more recent pooled analysis of 23 case-control studies, the *MTHFR* C677T variant rs1801133 was found to be protective against prostate cancer (OR per each additional T allele, 0.83; 95% CI 0.70-1.02) albeit with a wide confidence interval that included possible adverse effects[47]. The suggested protective effect of the folate reducing 677T allele runs counter to our MR study whereby each additional C allele (the folate increasing allele) confers a protective effect of similar magnitude (OR 0.87, see Figure 1A).

### Colorectal cancer

In our study, colorectal cancer was the only site-specific cancer to suggest increased risk with increasing serum folate, albeit with a wide confidence interval that included a possible protective effect. A 2018 systematic review[48] of RCTs reported little evidence of association between folic acid supplementation and colorectal cancer risk (OR 1.07; 95%CI 0.86-1.14) with effect direction in line with our MR analysis. For observational studies, the WCRF-CUP reported little evidence for association between dietary folate and colorectal cancer[43]. A large, recent meta-analysis (24 816 cases) reported reduced colorectal cancer risk when comparing highest (median, > 441 μg/day) with lowest (median, 212 μg/day) folate intake (RR 0.88; 95% CI 0.81-0.95)[16]. Using genetic studies, a meta-analysis of 67 studies reported strong evidence of association between the *MTHFR* 677TT genotype (which results in lower serum folate) and lower colorectal cancer risk under conditions of high folate intake[49].

### Ovarian cancer

Studies exploring the relationships between folate and risk of ovarian cancer are few. The ovarian WCRF-CUP, which was last updated in 2013, reported no evidence of associations between dietary folate or total folate and ovarian cancer in a dose-response meta-analysis (1158 cases) (RR 0.96; 95% CI 0.88-1.05)[43]. These results are in line with our weakly protective MR results.

### Lung cancer

In line with our MR results, the WCRF-CUP reported no significant associations between dietary folate intake and lung cancer; however, the report does report a possible U-shaped relationship (P-value < 0.01)[43]. Likewise, little evidence of association was observed for serum folate, though direction of effect was concordant with dietary intake and our MR study. Other recent meta-analyses of prospective cohort studies and RCTs also report little evidence of associations between folate intake (RR 0.99; 95% CI 0.97-1.01) and folic acid supplementation (RR 1.00; 95% CI 0.84-1.21) respectively[50,51].

### Malignant melanoma

Epidemiological studies relating folate and malignant melanoma risk are limited. An inverse relationship was reported for a meta-analysis of three RCTs evaluating treatment with combined supplements, including folic acid and risk for malignant melanomas (RR 0.47; 95% CI 0.23-0.94) in line with our current MR study. However, the sample size was very small (38 malignant melanoma cases)[52]. More recently, a meta-analysis of prospective cohorts reported a modest increased risk of malignant melanoma (1328 cases over a 26-year follow-up) for dietary folate intake (HR 1.36; 95% CI 1.13-1.64), though this did not replicate for total folate intake[53].

### Pan-cancer

Most studies published to date have focused on site-specific cancers. In this study, we have performed a genome-wide association analysis for pan-cancers. This allowed us to appraise the impact of folate on cancer risk across all sites in the general population. Proposed mechanisms for the formation of cancer via folate stems from the effects of perturbation of the one-carbon metabolism pathway effects of methylation and DNA repair and synthesis mechanisms which are common to the pathogenesis of many cancers[54]. However, we found little evidence of a causal association with pan-cancer, although the effect estimate was protective and of similar magnitude to those of the site-specific cancers. Likewise, a recent pooled analysis of RCTs showed that folic acid supplements had little effect on the risk of total cancer incidence (RR 1.06; 95% CI 0.99-1.13)[18]. The number of cancer cases was modest (3713 cases) and the mean follow-up time for included studies was five years, limiting conclusions of long-term impacts of folic acid supplementation.

In cancer treatment, antifolates are key compounds which inhibit enzymes in the folate metabolic pathway disrupting tumour growth and progression[55]. Due to the high proliferation of cells and demand for DNA, increasing levels of folate may promote the growth of precursor or established tumours in animal models[17,56]. Conversely, in normal tissues, insufficient folate levels may impair DNA replication and repair providing possible mechanisms for the initiation of cancer through gene mutation and chromosomal aberrations. Indeed, administration of folate has shown to reverse these effects[57]. This may support the general findings of protective effect estimates as reported for the multiple cancers within this MR study and in previously published studies. The role of folate in cancer treatment strategies, as well as the proposed mechanisms for carcinogenesis and progression, suggests that potential associations between folate and cancer may not be in terms of risk per se, but rather in progression and survival.

### Strengths and limitations

This study’s major strength is the use of two-sample MR, which is less prone to biases from confounding, reverse causation and measurement error that is seen in observational studies using directly measured phenotypes. Robust instruments for MR will have a biologically plausible relationship to the exposure (though this is not mandatory), i.e. located at or near genes with established pathways relevant to the exposure. The lead GWAS SNP rs1801133 (C667T;A222V) resides within the Methylenetetrahydrofolate reductase (*MTHFR*) gene, reduces *MTHFR* activity and increases it thermolability, in turn lowering serum folate levels, particularly in individuals with low dietary folate intake[58]. This satisfies the criteria of biological plausibility and thus strengthens the robustness of our serum folate instrument.

We also have several limitations that impact our interpretation of findings. We were unable to extend our analysis to allow for stratified analyses by factors of interest such as alcohol intake, BMI, sex, age, menopausal status and smoking. Our causal estimators assumed a linear relationship, and we were also unable to test for deviations from this. Several methods have recently been developed to explore non-linear relationships within an MR framework; however, these approaches are underpowered and require access to individual-level data[59].

We had greater than 90% power to detect our reported MR ORs for breast, prostate, ovarian, lung and colorectal cancer in the consortia GWAS datasets. However, we had lower power for the cancers appraised using UKBB. Where possible, we performed meta-analysis within each site-specific cancer; which increases statistical power; but we may still be underpowered to detect clinically relevant but smaller magnitude effects. Furthermore, statistical power in MR is dependent on the proportion of variance in the exposure variable explained by the genetic variants. The three SNPs within our MR were in weak LD with each other; therefore, it is likely that the variance explained is lower than that of the sum of the three SNPs (R^2^ 5%). Further work to identify additional SNPs robustly associated with serum folate in larger GWAS and meta-analysis will go some way to improving statistical power.

The SNPs included in our instrument were found to be associated with vascular traits, and cell and platelet measures in MR-Base PheWAS. Previous studies have reported associations between folate and vascular traits[60] as well as between vascular traits and cancer risk[61,62]. The serum folate instruments used in our MR are located within or near the *MTHFR* gene, which is directly involved in the one-carbon metabolism pathway. This suggests that the associations with vascular traits could reflect the downstream effects of folate on cancer rather than pleiotropic effects. Though there is no compelling evidence, we cannot rule out the possibility of associations with potential confounders or the presence of pleiotropy.

This study was conducted in European populations and so may not be generalisable to other populations. Both the Republic of Ireland and the UK voluntarily fortify cereals, which may help reduce the prevalence of folate-deficient individuals. Further work may be needed to explore causal effects in populations in which both samples (sample one and sample two) included in the MR analyses derive from countries with no such voluntary or mandatory fortification of food items.

### Conclusions

We found little evidence of causal associations between genetically elevated serum folate levels and the risk of pan-cancer, and cancer of the breast, prostate, ovaries, lung, colorectum; and malignant melanoma. Further work is needed to replicate our findings, strengthen the folate instrumental variable, and explore causal associations in the risk of cancer subtypes. In combination with existing literature, our results provide some evidence that increasing levels of circulating folate through widespread supplementation or deregulation of fortification of foods with folic acid is unlikely to lead to moderate unintended population-wide increase in cancer risk.

## Funding

This work was supported by a grant awarded to SJL by the World Cancer Research Fund International [WCRF 2015/1421], a Cancer Research UK program grant [C18281/A19169] and the National Institute for Health Research (NIHR) Bristol Biomedical Research Centre. PH is supported by CRUK Population Research Postdoctoral Fellowship [C52724/A20138]. KKT was supported by the World Cancer Research Fund (WCRF UK), as part of the World Cancer Research Fund International grant programme [WCRF 2014/1180].

The funders played no role in the design, implementation, analysis, or interpretation of the data in this study.

## Acknowledgements

The authors would like to thank the participants of the individual studies contributing to BCAC, PRACTICAL, OCAC, ILCCO, GECCO-CORECT-CCFR, and UKBB for their participation in these studies along with the principal investigators for generating the data and for making these data available or available within the public domain.

The authors would also like to thank Professor Barry Shane and Dr. Faith Pangilinan; authors of the study from which we defined our genetic instrument; for responding to our queries and providing additional information on data used within this manuscript.

Finally, we would like to thank our colleagues Dr. James Yarmolinsky, Dr. Caroline Bull, Dr. Vanessa Tan, Dr. Tom Dudding, and Dr. Timothy Robinson for their contributions towards defining cancer outcomes in UKBB.

## Contributors

SJL, RMM and KB conceived and designed the study. KB conducted the analyses. KB wrote the manuscript with input from all authors. Correspondence and material requests should be addressed to KB.

## Declaration of interests

None

## Notes

### Competing Interest Statement

The authors have declared no competing interest.

### Summary of Updates

Figure 1 has been revised. Supplemental files have been updated.

