## Supplementary Tables for "Mendelian randomisation study exploring the associations of serum folate with pan and site-specific cancers"

*Table S1 UK Biobank cancer registry case and control definitions*

|  | **Pan-cancer** | **Breast** | **Prostate** | **Ovarian** | **Lung** | **Colorectal** | **Malignant Melanoma** |
| --- | --- | --- | --- | --- | --- | --- | --- |
| **Cases** | | | | | | | |
| *ICD9 codes* | 140.0-208.9 | 174.0-174.9 | 185 | 183 | 162.2-162.5; 162.8-162.9 | 153.0-153.9 | 172.0-172.9 |
| *ICD10 codes* | All ICD10:C codes | C50.0-C50.9 | C61 | C56 | C34.0-C34.3; C34.8-C34.9 | C18.0-C18.9; C19; C20 | C43.0-C43.9 |
| *Inclusions* | Behaviour of tumour: "Malignant, primary site", "Malignant, microinvasive", "Malignant, metastatic site", "Malignant, uncertain whether primary or metastatic site" | | | | | | |
| *Exclusions* | Behaviour of tumour: "Benign", "Uncertain whether benign or malignant", "Carcinoma in situ"; ICD10:D codes | | | | | | |
|  | ICD10:C44.0-C44.9; ICD9:173.0-173.9 | Males | Females | Males |  |  | ICD10:C44.0-C44.9; ICD9:173.0-173.9 |
| **Controls** | | | | | | | |
| *Inclusions* | All eligible participants who are not defined as cases | | | | | | |
|  | Males and females | Females only | Males only | Females only | Males and females | Males and females | Males and females |
| *Exclusions* | Any ICD9 cancer code (140.0-239.7); any ICD10:C code; ICD10:D code (in situ & benign neoplasms); self-report of cancer or any site-specific cancer | | | | | | |

*Table S2 Correlation between serum folate genetic variants*

| **SNP** | rs1801133_A_G | rs7545014_T_C | rs1999594_A_G |
| --- | --- | --- | --- |
| rs1801133_A_G | 1 | 0.05 | 0.24 |
| rs7545014_T_C | 0.22 | 1 | 0.44 |
| rs1999594_A_G | -0.49 | -0.67 | 1 |

*Upper triangle represents R^2^, lower triangle represents Pearson’s R for correlation*

Table S3 Study sample sizes and power calculations for pan-cancer and six site-specific cancers

|  | **Study** | **Cases** | **Controls** | **Sample size** | **% cases** | **OR (MR)** | **Power (MR)** | **Power at OR:** | | | | | |
| --- | --- | --- | --- | --- | --- | --- | --- | --- | --- | --- | --- | --- | --- |
| **Cancer type** |  |  |  |  |  |  |  | **1.05** | **1.1** | **1.2** | **1.3** | **1.4** | **1.5** |
| **Pan-cancers** | UKBB | 50643 | 372016 | 422659 | 11.98 | **0.88** | **1.00** | 0.65 | 1.00 | 1.00 | 1.00 | 1.00 | 1.00 |
| **Breast** | BCAC | 122977 | 105974 | 228951 | 53.71 | **0.94** | **0.93** | 0.74 | 1.00 | 1.00 | 1.00 | 1.00 | 1.00 |
|  | UKBB | 13879 | 198523 | 212402 | 6.53 | **0.83** | **0.99** | 0.24 | 0.71 | 1.00 | 1.00 | 1.00 | 1.00 |
| **Prostate** | PRACTICAL* | 79148 | 61106 | 140254 | 56.43 | **0.86** | **1.00** | 0.52 | 0.98 | 1.00 | 1.00 | 1.00 | 1.00 |
|  | UKBB | 9132 | 173493 | 182625 | 5.00 | **0.89** | **0.63** | 0.18 | 0.54 | 0.98 | 1.00 | 1.00 | 1.00 |
| **Ovarian** | OCAC | 25509 | 40941 | 66450 | 38.39 | **0.86** | **0.98** | 0.28 | 0.77 | 1.00 | 1.00 | 1.00 | 1.00 |
|  | UKBB | 1218 | 198523 | 199741 | 0.61 | **0.67** | **0.72** | 0.07 | 0.12 | 0.34 | 0.64 | 0.87 | 0.97 |
| **Lung** | ILCCO | 11348 | 15861 | 27209 | 41.71 | **0.81** | **0.97** | 0.14 | 0.42 | 0.92 | 1.00 | 1.00 | 1.00 |
|  | UKBB | 2671 | 372016 | 374687 | 0.71 | **1.02** | **0.06** | 0.09 | 0.21 | 0.63 | 0.93 | 1.00 | 1.00 |
| **Colorectal** | GECCO* | 58221 | 67694 | 125915 | 46.24 | **1.18** | **1.00** | 0.49 | 0.97 | 1.00 | 1.00 | 1.00 | 1.00 |
|  | UKBB | 5657 | 372016 | 377673 | 1.50 | **1.45** | **1.00** | 0.67 | 0.39 | 0.91 | 1.00 | 1.00 | 1.00 |
| **Malignant melanoma** | UKBB | 3751 | 372016 | 375767 | 1.00 | **0.56** | **1.00** | 0.10 | 0.28 | 0.78 | 0.98 | 1.00 | 1.00 |

*OR (MR), odds ratio from MR analysis (IVW); Power (MR), power to detect the MR OR reported; Power (OR), power to detect a causal estimate at an alpha 0.05, R^2^ variance explained of 5% and the corresponding sample size and proportion of cases as indicated; ^*^ the PRACTICAL Consortium, CRUK, BPC3, CAPS and PEGASUS and the GECCO, CORECT and CCFR consortia.*

*Table S4 Wald ratio estimates for each SNP included in as an instrument for serum folate*

| Cancer type | Study | rs1801133 | | | rs1999594 | | | rs7545014 | | |
| --- | --- | --- | --- | --- | --- | --- | --- | --- | --- | --- |
|  |  | OR | SE | P Value | OR | SE | P Value | OR | SE | P Value |
| **Pan-cancers** | UKBB | 0.833 | 0.115 | 0.110 | 0.812 | 0.136 | 0.124 | 0.951 | 0.134 | 0.709 |
| **Breast** | BCAC | 1.041 | 0.112 | 0.717 | 0.867 | 0.124 | 0.252 | 0.818 | 0.125 | 0.109 |
|  | UKBB | 0.761 | 0.212 | 0.198 | 0.536 | 0.250 | 0.013 | 0.911 | 0.248 | 0.707 |
| **Prostate** | PRACTICAL* | 0.781 | 0.144 | 0.086 | 0.751 | 0.162 | 0.077 | 0.986 | 0.163 | 0.932 |
|  | UKBB | 1.076 | 0.259 | 0.777 | 0.824 | 0.307 | 0.529 | 0.665 | 0.305 | 0.181 |
| **Ovarian** | OCAC | 0.744 | 0.230 | 0.198 | 0.914 | 0.265 | 0.733 | 1.068 | 0.265 | 0.805 |
|  | UKBB | 0.464 | 0.696 | 0.270 | 0.323 | 0.822 | 0.169 | 1.115 | 0.814 | 0.893 |
| **Lung** | ILCCO | 0.806 | 0.308 | 0.484 | 0.937 | 0.358 | 0.857 | 0.814 | 0.362 | 0.571 |
|  | UKBB | 0.688 | 0.469 | 0.426 | 0.987 | 0.556 | 0.981 | 1.816 | 0.551 | 0.278 |
| **Colorectal** | GECCO* | 1.185 | 0.150 | 0.259 | 0.771 | 0.176 | 0.140 | 1.149 | 0.175 | 0.426 |
|  | UKBB | 1.363 | 0.324 | 0.339 | 1.190 | 0.383 | 0.650 | 1.570 | 0.380 | 0.235 |
| **Skin** | UKBB | 0.498 | 0.397 | 0.079 | 0.557 | 0.470 | 0.213 | 0.672 | 0.466 | 0.393 |

*OR, Wald MR odds ratio; SE, standard error; ORs represent the risk in cancer per SD increase in log_10_ serum folate; ^*^ the PRACTICAL Consortium, CRUK, BPC3, CAPS and PEGASUS and the GECCO, CORECT and CCFR consortia.*

*Table S5 Meta-analysis and Cochran’s Q Heterogeneity results for MR*

| **Cancer** | **OR** | **LCI** | **UCI** | **Pvalue** | **Q** | **Het P** |
| --- | --- | --- | --- | --- | --- | --- |
| **Breast** | 0.93 | 0.78 | 1.11 | 0.42 | 0.12 | 0.72 |
| **Prostate** | 0.87 | 0.71 | 1.06 | 0.17 | 0.02 | 0.89 |
| **Ovarian** | 0.84 | 0.59 | 1.20 | 0.34 | 0.17 | 0.68 |
| **Lung** | 0.87 | 0.57 | 1.32 | 0.51 | 0.24 | 0.63 |

*OR, odds ratio; LCI, lower 95% CI; UCI, upper 95% CI; Q Cochran’s Q statistic; Het P, P value for heterogeneity between meta-analysis studies*

*Table S6 Genome-wide significant associations of serum folate SNPs and traits in MR-Base*

| **Associations in MR-Base PHEWAS** | **P value** | **Sample Size** | **Cohort** | **Published** |
| --- | --- | --- | --- | --- |
| ***rs1801133*** |  |  |  |  |
| Diastolic blood pressure, automated reading | 2.20E-14 | 436324 | UKBB (MRC-IEU) | Unpublished |
| Diastolic blood pressure, automated reading | 4.06E-13 | 317756 | UKBB (Neal Lab) | Unpublished |
| Mean corpuscular hemoglobin | 1.17E-08 | 172332 | UK Biobank+INTERVAL+UK BiLEVE | <https://doi.org/10.1016/j.cell.2016.10.042> |
| Birth weight of first child | 4.10E-08 | 200272 | UKBB (MRC-IEU) | Unpublished |
| Mean corpuscular volume | 5.23E-08 | 172433 | UK Biobank+INTERVAL+UK BiLEVE | <https://doi.org/10.1016/j.cell.2016.10.042> |
| Plateletcrit | 1.74E-06 | 164339 | UK Biobank+INTERVAL+UK BiLEVE | <https://doi.org/10.1016/j.cell.2016.10.042> |
| Red cell distribution width | 2.33E-06 | 171529 | UK Biobank+INTERVAL+UK BiLEVE | <https://doi.org/10.1016/j.cell.2016.10.042> |
| Medication for cholesterol, blood pressure or diabetes: Blood pressure medication | 3.70E-06 | 209638 | UKBB (MRC-IEU) | Unpublished |
| Vascular/heart problems diagnosed by doctor: High blood pressure | 4.20E-06 | 461880 | UKBB (MRC-IEU) | Unpublished |
| Vascular/heart problems diagnosed by doctor: High blood pressure | 4.70E-06 | 336683 | UKBB (Neal Lab) | Unpublished |
| Non-cancer illness code, self-reported: hypertension | 4.90E-06 | 462933 | UKBB (MRC-IEU) | Unpublished |
| Hip circumference | 5.40E-06 | 462117 | UKBB (MRC-IEU) | Unpublished |
| Non-cancer illness code self-reported: hypertension | 9.64E-06 | 337159 | UKBB (Neal Lab) | Unpublished |
| ***rs7545014*** |  |  |  |  |
| Plateletcrit | 8.49E-11 | 164339 | UK Biobank+INTERVAL+UK BiLEVE | <https://doi.org/10.1016/j.cell.2016.10.042> |
| Platelet count | 1.63E-08 | 166066 | UK Biobank+INTERVAL+UK BiLEVE | <https://doi.org/10.1016/j.cell.2016.10.042> |
| Operative procedures - secondary OPCS: T29.1 Excision of umbilicus | 9.20E-06 | 463010 | UKBB (MRC-IEU) | Unpublished |
| ***rs1999594*** |  |  |  |  |
| Plateletcrit | 1.14E-15 | 164339 | UK Biobank+INTERVAL+UK BiLEVE | <https://doi.org/10.1016/j.cell.2016.10.042> |
| Platelet count | 6.36E-13 | 166066 | UK Biobank+INTERVAL+UK BiLEVE | <https://doi.org/10.1016/j.cell.2016.10.042> |
| Diastolic blood pressure, automated reading | 7.20E-06 | 436424 | UKBB (MRC-IEU) | Unpublished |

*Look-up performed in MR-Base (March 2019); cohorts UKBB (MRC-IEU) and (Neal Lab) indicates the GWAS performed on all traits in UKBB using automated pipelines. These results are not published but are publicly available to aid in identification of associations. This is on the proviso that researchers further explore these associations in more depth.*

Table S6 summarises the top associations detected between the serum folate SNPs with traits within the UKBB, GWAS catalog and curated datasets accessed via MR-Base. Only trait associations with P values <1e-05 are shown

*Table S7 Selected folate-cancer association estimates from observational studies*

| Cancer | Method | Risk Ratio | 95% CI Lower | 95% CI Upper | Units | Cases | Studies | Comment | Source |
| --- | --- | --- | --- | --- | --- | --- | --- | --- | --- |
| Breast | Total Folate Intake | 1.01 | 0.96 | 1.06 | 100 μg/day | 6094 | 3 | (I^2^ 70%, pval 0.04); CUP updated Jan 2017 | WCRF-CUP; https://www.wcrf.org/dietandcancer/breast-cancer |
| Breast | Dietary Folate Intake | 0.98 | 0.96 | 1.02 | 100 μg/day | 19251 | 6 | (I^2^ 0%, pval 0.81); CUP updated Jan 2017 |  |
| Prostate | Serum Folate Levels | 1.01 | 1.00 | 1.02 | 5 nmol/L | 5938 | 7 | (I^2^ 49%, pval 0.07); CUP updated Sep 2014 | WCRF-CUP; https://www.wcrf.org/dietandcancer/prostate-cancer |
| Ovarian | Total Folate Intake | 1.00* | 0.94 | 1.06 | 100 μg/day | 908 | 3 | (I^2^ 0%, pval 0.53); CUP updated Dec 2013 | WCRF-CUP; https://www.wcrf.org/dietandcancer/ovarian-cancer |
| Ovarian | Dietary Folate Intake | 0.92 | 0.77 | 1.10 | 100 μg/day | 1158 | 4 | (I^2^ 35%, pval 0.20); CUP updated Dec 2013 |  |
| Lung | Dietary Folate Intake | 0.99 | 0.95 | 1.02 | 100 μg/day | 4900 | 9 | (I^2^ 44%, pval 0.08); CUP updated Dec 2016 | WCRF-CUP; https://www.wcrf.org/dietandcancer/lung-cancer |
| Lung | Serum Folate Levels | 0.94 | 0.85 | 1.06 | 5 nmol/L | NA | 4 | From 2005 SLR (mentioned in CUP) |  |
| Colorectal | Dietary Folate Intake | 0.99 | 0.96 | 1.02 | 100 μg/day | 6986 | 10 | (I^2^ 31%, pval 0.16); CUP updated Sep 2017 | WCRF-CUP; https://www.wcrf.org/dietandcancer/colorectal-cancer |
| Colorectal | Total Folate Intake | 0.99 | 0.98 | 1.00 | 100 μg/day | 4633 | 8 | (I^2^ 0%, pval 0.92); CUP updated Sep 2017 |  |
| Colorectal | Serum Folate Levels | 0.99 | 0.98 | 1.01 | 5 nmol/L | 4261 | 12 | (I^2^ 4%, pval 0.41); CUP updated Sep 2017 |  |

*Estimates were used in Figure 2 of main paper; FFQ, food frequency questionnaire; RR, risk ratio; CI confidence intervals*
