## Supplementary Figures for "Mendelian randomisation study exploring the associations of serum folate with pan and site-specific cancers"

**Figure S1 – Scatter plots of SNP-folate and SNP-cancer effects**

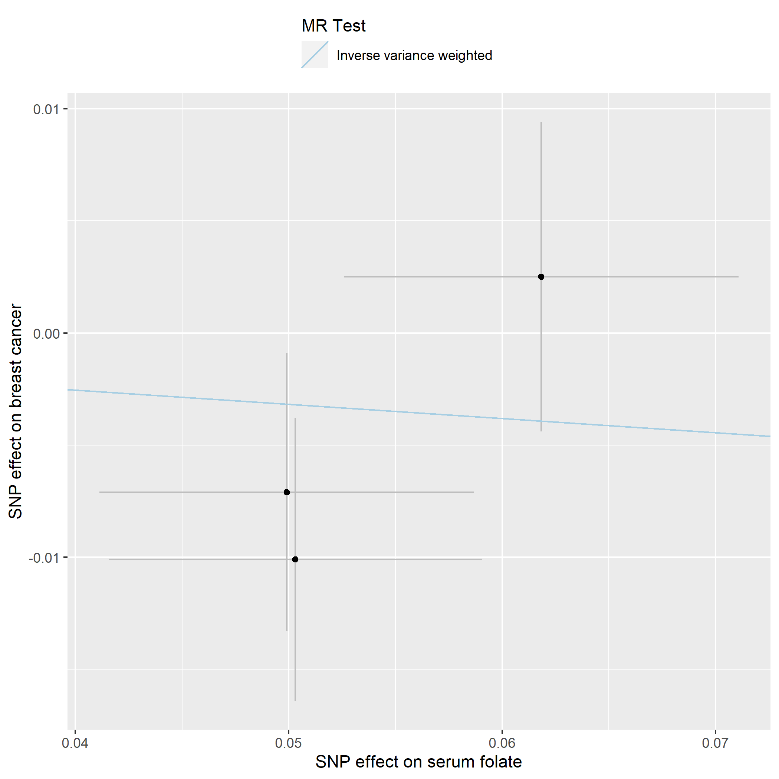

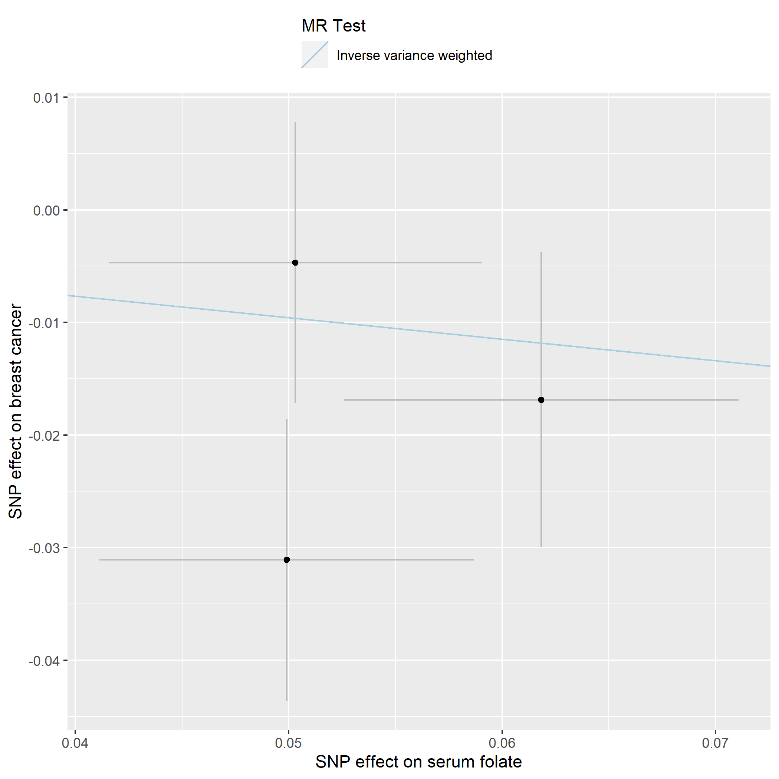

B

A

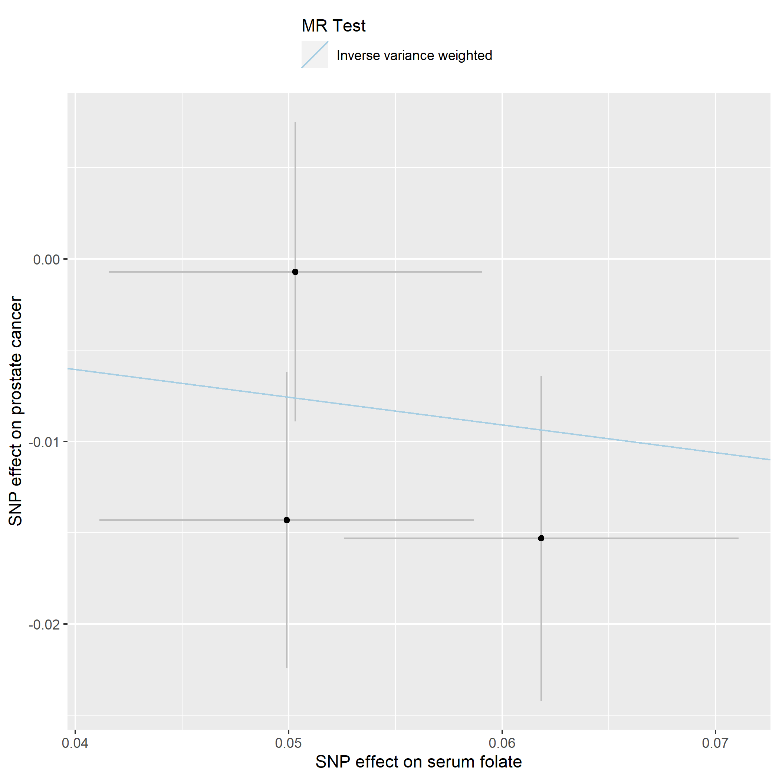

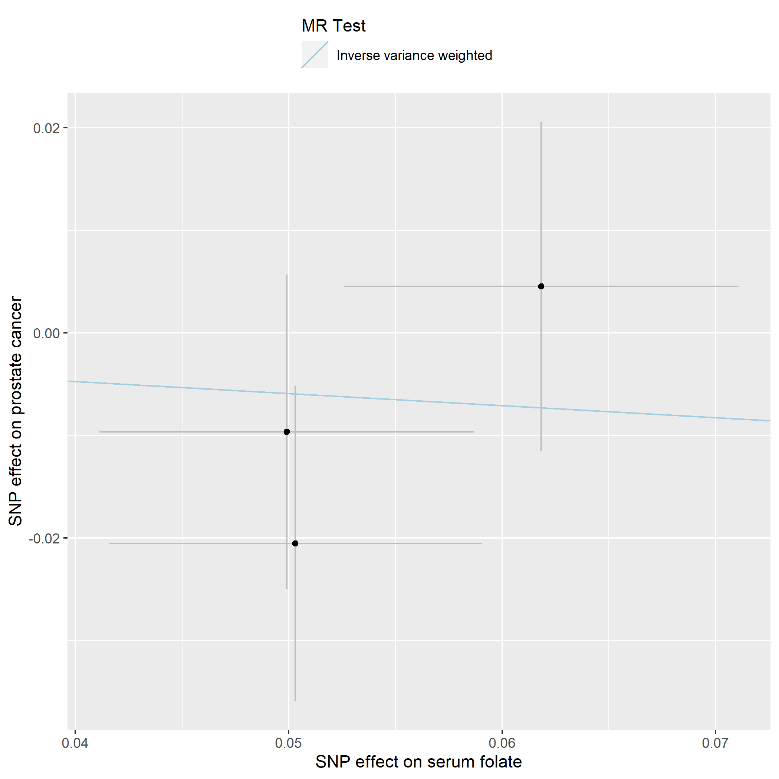

D

C

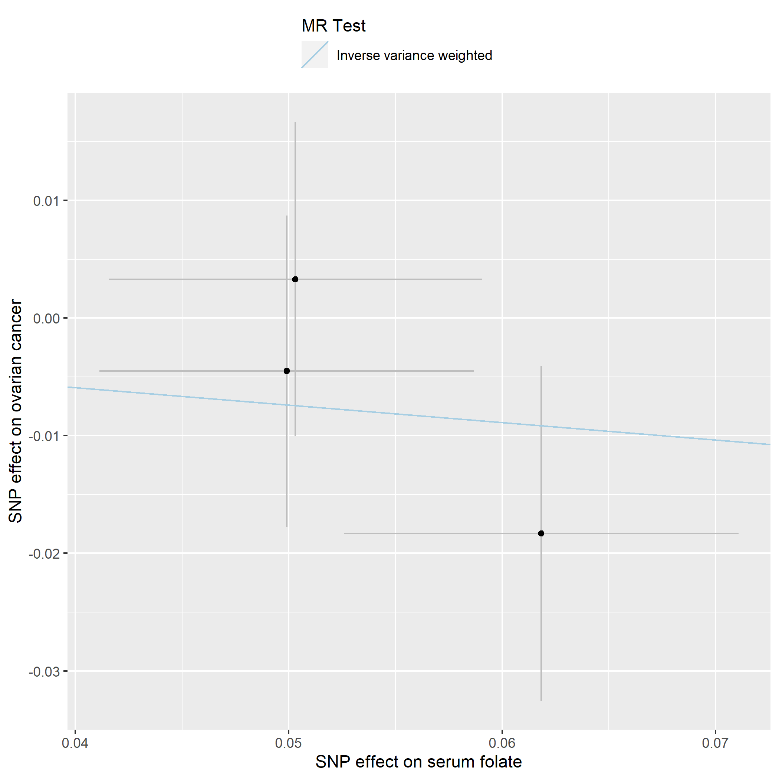

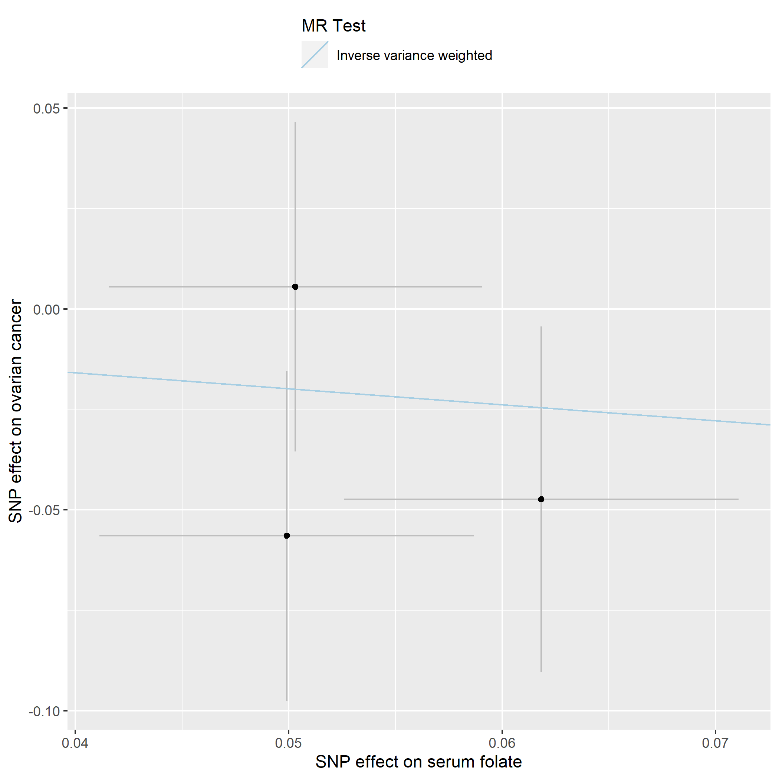

F

E

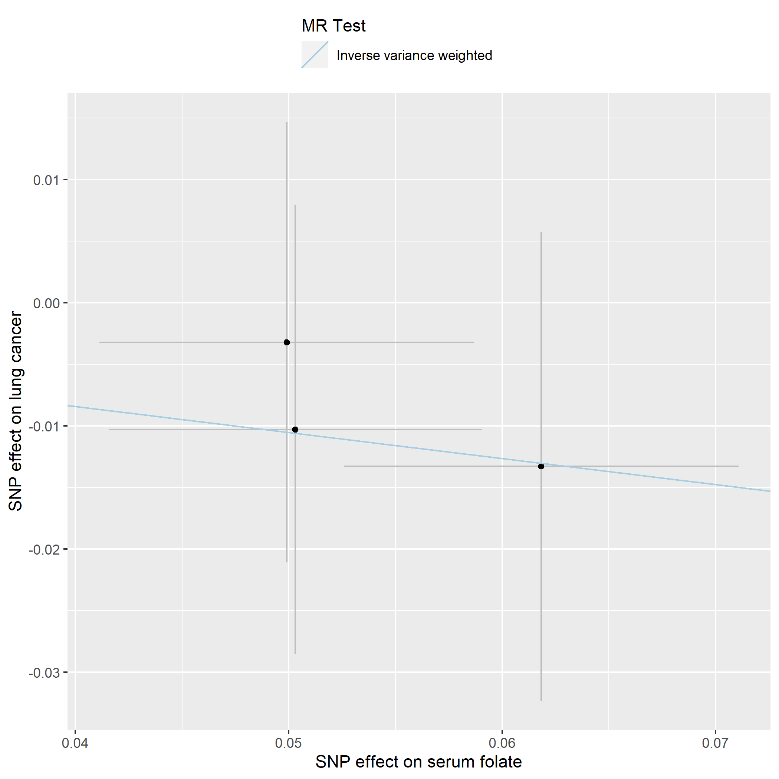

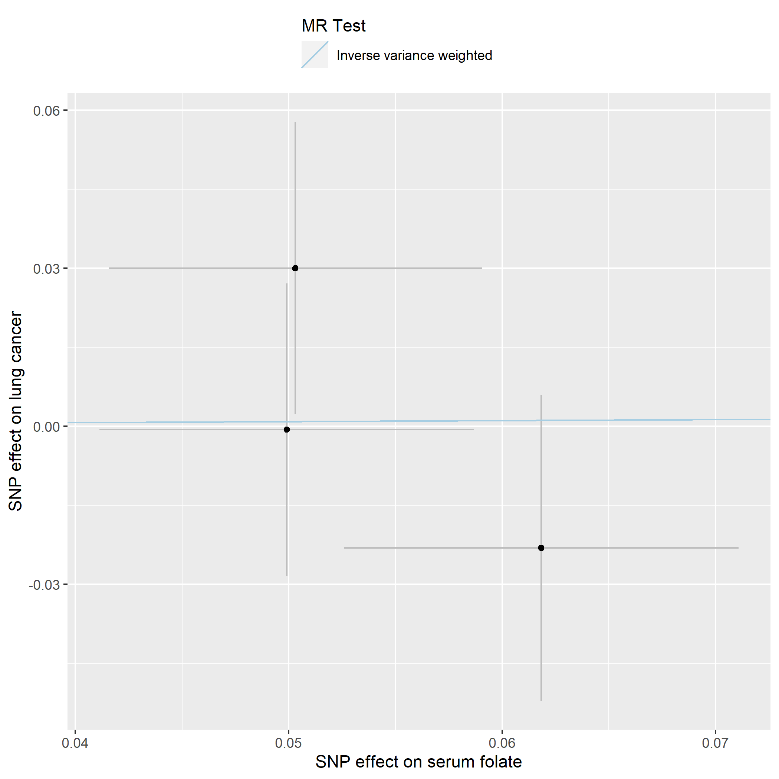

H

G

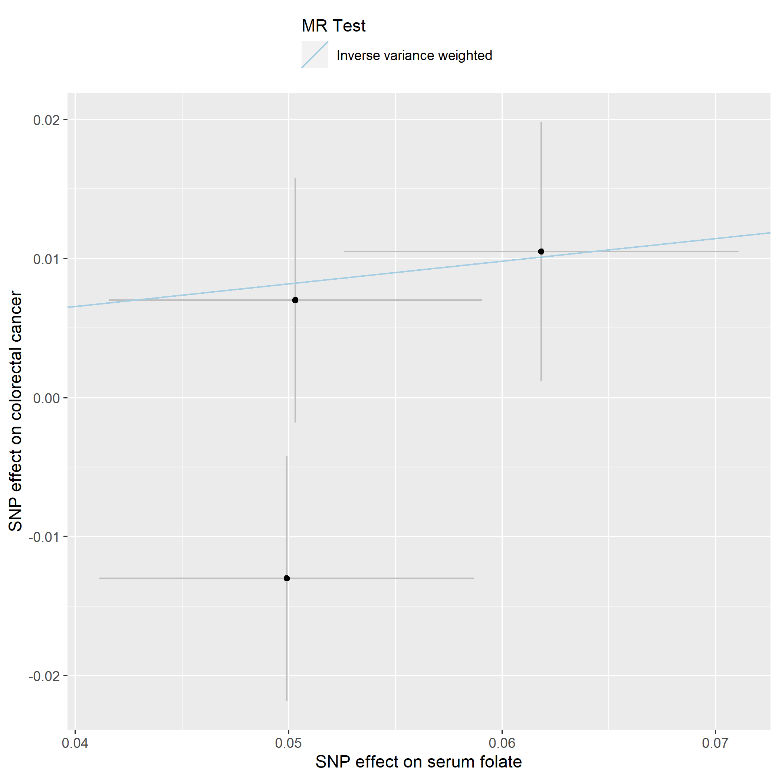

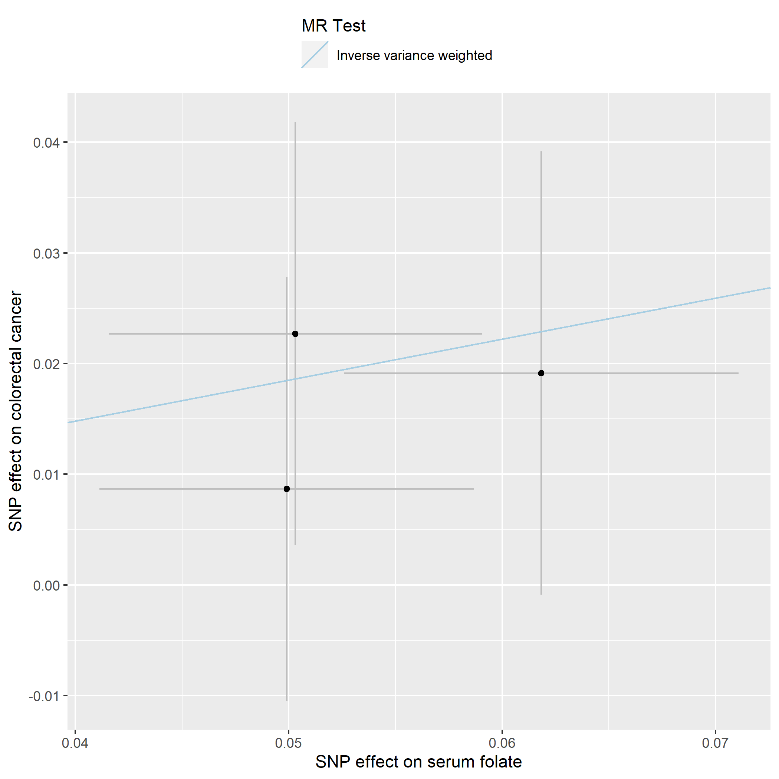

J

I

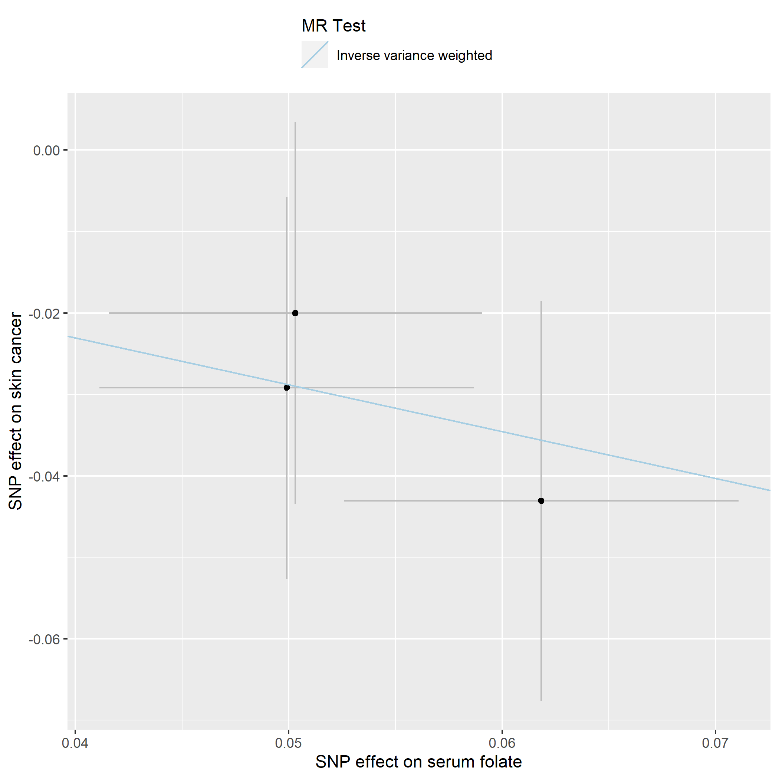

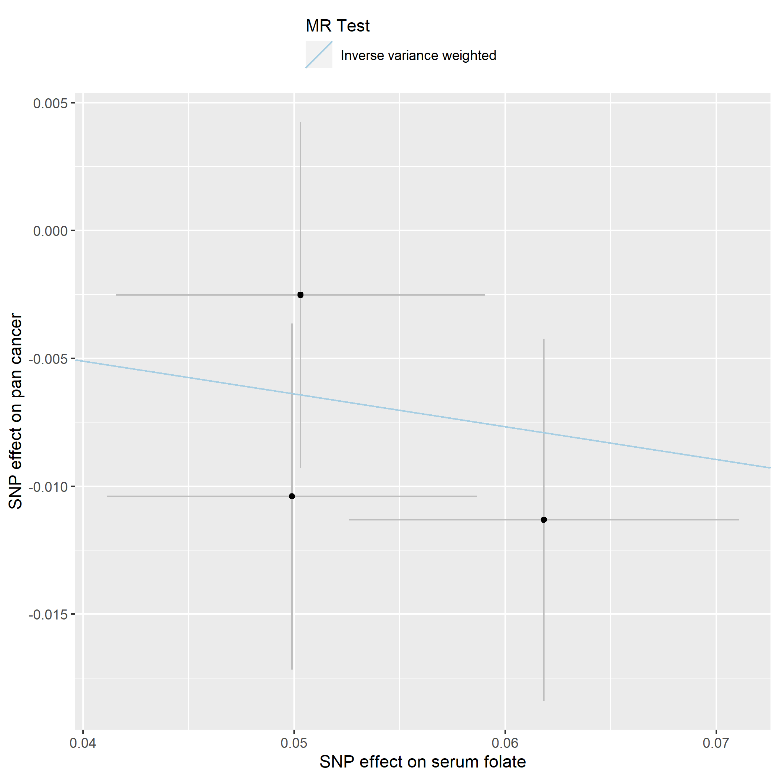

L

K

*A - BCAC: breast cancer; B - UKBB: breast cancer; C - PRACTICAL Consortium, CRUK, BPC3, CAPS and PEGASUS: prostate cancer; D - UKBB: prostate cancer; E - OCAC: ovarian cancer; F - UKBB: ovarian cancer; G - ILCCO: lung cancer; H - UKBB: lung cancer; I - GECCO, CORECT and CCFR: colorectal cancer; J - UKBB: colorectal cancer; K - UKBB: malignant melanoma; L - UKBB: pan-cancer.*

**Figure S2 – Forest plot of ORs in leave-one-out analysis**

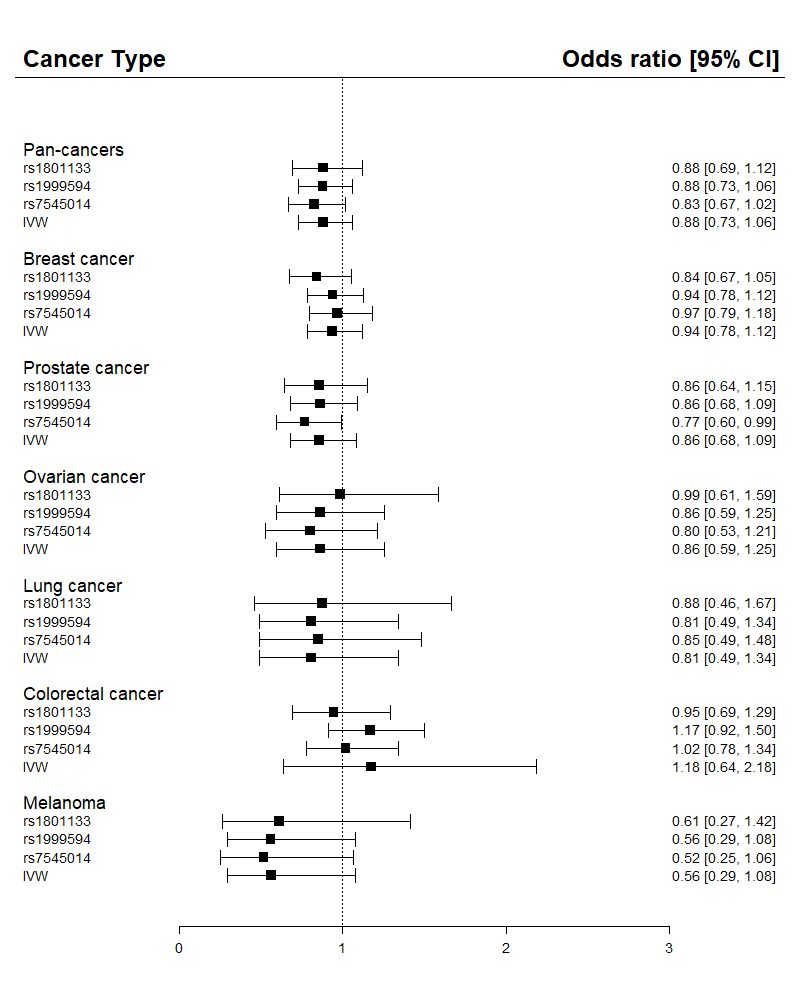

*IVW is the inverse variance weighted MR effect estimates of the analyses in the consortia GWAS for breast cancer (BCAC), prostate cancer (PRACTICAL Consortium, CRUK, BPC3, CAPS and PEGASUS), ovarian cancer (OCAC), colorectal cancer (GECCO, CORECT and CCFR), and lung cancer (ILCCO). IVW estimates are shown for UKBB analyses of pan-cancer and malignant melanoma.*
