## Supplementary Information for "Mendelian randomisation study exploring the associations of serum folate with pan and site-specific cancers"

Supplementary Material

### Cohort Information, funding and acknowledgements

The UK Biobank (UKBB)

We performed GWAS using data from the UK Biobank. UK Biobank is a population-based health research resource consisting of approximately 500,000 people, aged between 38 years and 73 years, who were recruited between the years 2006 and 2010 from across the UK[1]. Particularly focused on identifying determinants of human diseases in middle-aged and older individuals, participants provided a range of information (such as demographics, health status, lifestyle measures, cognitive testing, personality self-report, and physical and mental health measures) via questionnaires and interviews; anthropometric measures, BP readings and samples of blood, urine and saliva were also taken (data available at www.ukbiobank.ac.uk). A full description of the study design, participants and quality control (QC) methods have been described in detail previously[2]. UK Biobank received ethical approval from the Research Ethics Committee (REC reference for UK Biobank is 11/NW/0382).

Quality control filtering of the UK Biobank genetic data was conducted by R.Mitchell, G.Hemani, T.Dudding, L.Corbin, S.Harrison, L.Paternoster as described in the published protocol (doi:10.5523/bris.1ovaau5sxunp2cv8rcy88688v).

The MRC IEU UK Biobank GWAS pipeline was developed by B.Elsworth, R.Mitchell, C.Raistrick, L.Paternoster, G.Hemani, T.Gaunt (doi:10.5523/bris.pnoat8cxo0u52p6ynfaekeigi.).

UKBB data was accessed under the application number 15825 (Dr. Philip Haycock).

**Cancer cases and control definitions**

Table S1 gives the case and control definitions for each of the site-specific and pan-cancer. Briefly, The UK Biobank extracted records of all cancers including their subtype occurring either before or after participant enrolment using the UK cancer registry. Cancers were recorded according to the International Classification of Diseases (ICD9, ICD10). In addition, participants also self-reported cancers. Both cancer registry and self-report of cancer were taken at multiple follow-up timepoints. Cancer registry data was updated to April 2019.

Site-specific cancer cases were defined as a diagnosis captured by the registry according to relevant ICD9 and 10 codes (see Table S1). Cases were excluded if the record for “cancer tumour behaviour” were any of the following: benign, uncertain whether benign or malignant, or carcinoma in situ. For pan-cancer, cases were defined as individuals who have reported diagnosis of any of the ICD9 and ICD10 malignant cancer codes except for non-melanoma skin cancers. The list of cancer codes is given in Table S1. Cases were again excluded according to the cancer tumour behaviour definitions as given above. For both pan-cancers and site-specific cancers controls were defined as individuals who do not have any report of diagnosis of any cancers or carcinomas *in situ,* or undefined neoplasm. Controls were further defined as also having no self-report of cancers.

Breast Cancer Association Consortium (BCAC)

We used published genome-wide association data reported by the Breast Cancer Association Consortium (BCAC; <http://bcac.ccge.medschl.cam.ac.uk/bcacdata/oncoarray/gwas-icogs-and-oncoarray-summary-results/>). SNP-cancer estimates were reported for meta-analysis including BCAC, Discovery, Biology and Risk of Inherited Variants in Breast Cancer Consortium (DRIVE), Collaborative Oncological Gene-environment Study (iCOGS), and 11 other breast cancer genome-wide association studies pooling 122,977 cases and 105,974 controls for overall breast cancer. Participants were of European ancestry and details on the genotyping protocol, GWAS and meta-analysis have been previously described[3].

**Funding and acknowledgements**

The breast cancer genome-wide association analyses were supported by the Government of Canada through Genome Canada and the Canadian Institutes of Health Research, the ‘Ministère de l’Économie, de la Science et de l’Innovation du Québec’ through Genome Québec and grant PSR-SIIRI-701, The National Institutes of Health (U19 CA148065, X01HG007492), Cancer Research UK (C1287/A10118, C1287/A16563, C1287/A10710) and The European Union (HEALTH-F2-2009-223175 and H2020 633784 and 634935). All studies and funders are listed in Michailidou et al (2017)[3].

Prostate Cancer Association Group to Investigate Cancer Associated Alterations in the Genome (PRACTICAL)

We used published genome-wide association data reported by the Prostate Cancer Association Group to Investigate Cancer-Associated Alterations in the Genome (PRACTICAL). Summary data were derived from a meta-analysis of GWAS of overall prostate cancer in Europeans resulting in 79,148 cases and 61,106 controls. Details on the genotyping protocol, GWAS and meta-analysis have been previously described[4].

**Authors, funding and acknowledgements**

*PIs from the PRACTICAL (http://practical.icr.ac.uk/), CRUK, BPC3, CAPS, PEGASUS consortia:*

Rosalind A. Eeles1,2, Brian E. Henderson3**, Christopher A. Haiman3, ZSofia Kote-Jarai1, Fredrick R. Schumacher4,5, Sara Benlloch6,1, Ali Amin Al Olama6,9, Kenneth Muir7,8, Sonja I. Berndt10, David V. Conti3, Fredrik Wiklund11, Stephen Chanock10, Susan M. Gapstur12, Victoria L. Stevens12, Catherine M. Tangen13, Jyotsna Batra14,16, Judith Clements15,16, APCB BioResource15, Henrik Gronberg11, Nora Pashayan17,18, Johanna Schleutker19,20, Demetrius Albanes21, Stephanie Weinstein21, Alicja Wolk22, Catharine West24, Lorelei Mucci25, Géraldine Cancel-Tassin26,27, Stella Koutros10, Karina Dalsgaard Sorensen28,29, Eli Marie Grindedal30, David E. Neal31,32,33, Freddie C. Hamdy34,35, Jenny L. Donovan36, Ruth C. Travis37, Robert J. Hamilton38,39, Sue Ann Ingles3, Barry S. Rosenstein40, Yong-Jie Lu41, Graham G. Giles42,43,44, Adam S. Kibel45, Ana Vega46,47,48, Manolis Kogevinas49,50,51,52, Kathryn L. Penney53, Jong Y. Park54, Janet L. Stanford55,56, Cezary Cybulski57, Børge G. Nordestgaard58,59, Sune F. Nielsen58,59, Hermann Brenner60,61,62, Christiane Maier63, Jeri Kim64, Esther M. John65, Manuel R. Teixeira66,67, Susan L. Neuhausen68, Kim De Ruyck69, Azad Razack70, Lisa F. Newcomb55,71, Davor Lessel72, Radka Kaneva73, Nawaid Usmani74,75, Frank Claessens76, Paul A. Townsend77, Manuela Gago-Dominguez78,79, Monique J. Roobol80, Florence Menegaux81, Kay-Tee Khaw82, Lisa Cannon-Albright83,84, Hardev Pandha85, Stephen N. Thibodeau86, David J Hunter87, Peter Kraft87, William J. Blot88,89, Elio Riboli90,

**In memorium

1	The Institute of Cancer Research, London, UK.

2	Royal Marsden NHS Foundation Trust, London, UK.

3	Department of Preventive Medicine, Keck School of Medicine, University of Southern California/Norris Comprehensive Cancer Center, Los Angeles, CA 90015, USA

4	Department of Population and Quantitative Health Sciences, Case Western Reserve University, Cleveland, OH 44106-7219, USA

5	Seidman Cancer Center, University Hospitals, Cleveland, OH 44106, USA.

6	Centre for Cancer Genetic Epidemiology, Department of Public Health and Primary Care, University of Cambridge, Strangeways Research Laboratory, Cambridge, UK

7	Division of Population Health, Health Services Research and Primary Care, University of Manchester, Oxford Road, Manchester, M13 9PL, UK

8	Warwick Medical School, University of Warwick, Coventry, UK.

9	University of Cambridge, Department of Clinical Neurosciences, Stroke Research Group, R3, Box 83, Cambridge Biomedical Campus, Cambridge CB2 0QQ, UK

10	Division of Cancer Epidemiology and Genetics, National Cancer Institute, NIH, Bethesda, Maryland, 20892, USA

11	Department of Medical Epidemiology and Biostatistics, Karolinska Institute, Stockholm, Sweden.

12	Epidemiology Research Program, American Cancer Society, 250 Williams Street, Atlanta, GA 30303, USA

13	SWOG Statistical Center, Fred Hutchinson Cancer Research Center, Seattle, WA 98109, USA

14	Australian Prostate Cancer Research Centre-Qld, Institute of Health and Biomedical Innovation and School of Biomedical Sciences, Queensland University of Technology, Brisbane, QLD 4059, Australia

15	Australian Prostate Cancer Research Centre-Qld, Institute of Health and Biomedical Innovation and School of Biomedical Sciences, Queensland University of Technology, Brisbane QLD 4059, Australia

16	Translational Research Institute, Brisbane, Queensland 4102, Australia

17	University College London, Department of Applied Health Research, London, UK.

18	Centre for Cancer Genetic Epidemiology, Department of Oncology, University of Cambridge, Strangeways Laboratory, Cambridge, UK.

19	Institute of Biomedicine, Kiinamyllynkatu 10, FI-20014 University of Turku, Finland

20	Department of Medical Genetics, Genomics, Laboratory Division, Turku University Hospital, PO Box 52, 20521 Turku, Finland

21	Division of Cancer Epidemiology and Genetics, National Cancer Institute, NIH, Bethesda, MD 20892, USA

22	Division of Nutritional Epidemiology, Institute of Environmental Medicine, Karolinska Institutet, SE-171 77 Stockholm, Sweden

23	Department of Surgical Sciences, Uppsala University, Uppsala, Sweden

24	Division of Cancer Sciences, University of Manchester, Manchester Academic Health Science Centre, Radiotherapy Related Research, The Christie Hospital NHS Foundation Trust, Manchester, M13 9PL UK

25	Department of Epidemiology, Harvard T.H. Chan School of Pubic Health, Boston, MA, USA.

26	CeRePP, Tenon Hospital, Paris, France

27	Sorbonne Universite, GRC n°5 ONCOTYPE-URO, AP-HP, Tenon Hospital, 4 rue de la Chine, F-75020 Paris, France

28	Department of Molecular Medicine, Aarhus University Hospital, Palle Juul-Jensen Boulevard 99, 8200 Aarhus N, Denmark

29	Department of Clinical Medicine, Aarhus University, DK-8200 Aarhus N

30	Department of Medical Genetics, Oslo University Hospital, Norway.

31	Nuffield Department of Surgical Sciences, University of Oxford, Room 6603, Level 6, John Radcliffe Hospital, Headley Way, Headington, Oxford, OX3 9DU, UK

32	University of Cambridge, Department of Oncology, Addenbrooke's Hospital, Cambridge, UK.

33	Cancer Research UK Cambridge Research Institute, Li Ka Shing Centre, Cambridge, UK.

34	Nuffield Department of Surgical Sciences, University of Oxford, Oxford, OX1 2JD, UK

35	Faculty of Medical Science, University of Oxford, John Radcliffe Hospital, Oxford, UK

36	Population Health Sciences, Bristol Medical School, University of Bristol, BS8 2PS, UK

37	Cancer Epidemiology Unit, Nuffield Department of Population Health University of Oxford, Oxford, UK.

38	Dept. of Surgical Oncology, Princess Margaret Cancer Centre, Toronto, Canada.

39	Dept. of Surgery (Urology), University of Toronto, Canada

40	Department of Radiation Oncology and Department of Genetics and Genomic Sciences, Box 1236, Icahn School of Medicine at Mount Sinai, One Gustave L. Levy Place, New York, NY 10029, USA

41	Centre for Molecular Oncology, Barts Cancer Institute, Queen Mary University of London, John Vane Science Centre, Charterhouse Square, London, EC1M 6BQ, UK

42	Cancer Epidemiology Division, Cancer Council Victoria, 615 St Kilda Road, Melbourne, Victoria 3004, Australia.

43	Centre for Epidemiology and Biostatistics, Melbourne School of Population and Global Health, The University of Melbourne, Grattan Street, Parkville, VIC 3010, Australia

44	Precision Medicine, School of Clinical Sciences at Monash Health. Monash University. Clayton, Victoria, Australia, 3168

45	Division of Urologic Surgery, Brigham and Womens Hospital, 75 Francis Street, Boston, MA 02115, USA

46	Fundación Pública Galega de Medicina Xenómica, Santiago de Compostela, 15706, Spain.

47	Instituto de Investigación Sanitaria de Santiago de Compostela, Santiago De Compostela, 15706, Spain

48	Centro de Investigación en Red de Enfermedades Raras (CIBERER), Spain

49	ISGlobal, Barcelona, Spain.

50	IMIM (Hospital del Mar Research Institute), Barcelona, Spain.

51	Universitat Pompeu Fabra (UPF), Barcelona, Spain.

52	CIBER Epidemiologia y Salud Publica (CIBERESP), Madrid, Spain

53	Channing Division of Network Medicine, Department of Medicine, Brigham and Women's Hospital/Harvard Medical School, Boston, MA, USA.

54	Department of Cancer Epidemiology, Moffitt Cancer Center, 12902 Magnolia Drive, Tampa, FL 33612, USA

55	Division of Public Health Sciences, Fred Hutchinson Cancer Research Center, Seattle, Washington, 98109-1024, USA

56	Department of Epidemiology, School of Public Health, University of Washington, Seattle, Washington, USA.

57	International Hereditary Cancer Center, Department of Genetics and Pathology, Pomeranian Medical University, Szczecin, Poland.

58	Faculty of Health and Medical Sciences, University of Copenhagen, Denmark.

59	Department of Clinical Biochemistry, Herlev and Gentofte Hospital, Copenhagen University Hospital, Herlev, Denmark.

60	Division of Clinical Epidemiology and Aging Research, German Cancer Research Center (DKFZ), Heidelberg, Germany.

61	German Cancer Consortium (DKTK), German Cancer Research Center (DKFZ), Heidelberg, Germany.

62	Division of Preventive Oncology, German Cancer Research Center (DKFZ) and National Center for Tumor Diseases (NCT), Heidelberg, Germany.

63	Institute for Human Genetics, University Hospital Ulm, Ulm, Germany.

64	The University of Texas M. D. Anderson Cancer Center, Department of Genitourinary Medical Oncology, Houston, TX, USA.

65	Department of Medicine, Division of Oncology, Stanford Cancer Institute, Stanford University School of Medicine, Stanford, 780 Welch Road, CJ250C, CA 94304-5769

66	Department of Genetics, Portuguese Oncology Institute of Porto, Porto, Portugal.

67	Biomedical Sciences Institute (ICBAS), University of Porto, Porto, Portugal.

68	Department of Population Sciences, Beckman Research Institute of the City of Hope, Duarte, CA, USA.

69	Ghent University, Faculty of Medicine and Health Sciences, Basic Medical Sciences, Gent, Belgium.

70	Department of Surgery, Faculty of Medicine, University of Malaya, Kuala Lumpur, Malaysia.

71	Department of Urology, University of Washington, 1959 NE Pacific Street, Box 356510, Seattle, WA 98195, USA

72	Institute of Human Genetics, University Medical Center Hamburg-Eppendorf, Hamburg, Germany.

73	Molecular Medicine Center, Department of Medical Chemistry and Biochemistry, Medical University, Sofia, Bulgaria.

74	Department of Oncology, Cross Cancer Institute, University of Alberta, Edmonton, Alberta, Canada.

75	Division of Radiation Oncology, Cross Cancer Institute, Edmonton, Alberta, Canada.

76	Molecular Endocrinology Laboratory, Department of Cellular and Molecular Medicine, KU Leuven, Leuven, Belgium.

77	Division of Cancer Sciences, Manchester Cancer Research Centre, Faculty of Biology, Medicine and Health, Manchester Academic Health Science Centre, NIHR Manchester Biomedical Research Centre, Health Innovation Manchester, Univeristy of Manchester, UK.

78	Genomic Medicine Group, Galician Foundation of Genomic Medicine, Instituto de Investigacion Sanitaria de Santiago de Compostela (IDIS), Complejo Hospitalario Universitario de Santiago, Servicio Galego de Saúde, SERGAS, Santiago De Compostela, Spain.

79	University of California San Diego, Moores Cancer Center, La Jolla, CA, USA.

80	Department of Urology, Erasmus University Medical Center, Rotterdam, the Netherlands.

81	Cancer & Environment Group, Center for Research in Epidemiology and Population Health (CESP), INSERM, University Paris-Sud, University Paris-Saclay, Villejuif, France.

82	Clinical Gerontology Unit, University of Cambridge, Cambridge, UK.

83	Division of Genetic Epidemiology, Department of Medicine, University of Utah School of Medicine, Salt Lake City, Utah, USA.

84	George E. Wahlen Department of Veterans Affairs Medical Center, Salt Lake City, UT, USA.

85	The University of Surrey, Guildford, Surrey, UK.

86	Department of Laboratory Medicine and Pathology, Mayo Clinic, Rochester, MN, USA.

87	Program in Genetic Epidemiology and Statistical Genetics, Department of Epidemiology, Harvard T.H. Chan School of Public Health, Boston, MA, USA.

88	Division of Epidemiology, Department of Medicine, Vanderbilt University Medical Center, TN, USA.

89	International Epidemiology Institute, Rockville, MD, USA

90	Department of Epidemiology and Biostatistics, School of Public Health, Imperial College London, SW7 2AZ, UK

*CRUK and PRACTICAL consortium*

This work was supported by the Canadian Institutes of Health Research, European Commission's Seventh Framework Programme grant agreement n° 223175 (HEALTH-F2-2009-223175), Cancer Research UK Grants C5047/A7357, C1287/A10118, C1287/A16563, C5047/A3354, C5047/A10692, C16913/A6135, and The National Institute of Health (NIH) Cancer Post-Cancer GWAS initiative grant: No. 1 U19 CA 148537-01 (the GAME-ON initiative).

We would also like to thank the following for funding support: The Institute of Cancer Research and The Everyman Campaign, The Prostate Cancer Research Foundation, Prostate Research Campaign UK (now PCUK), The Orchid Cancer Appeal, Rosetrees Trust, The National Cancer Research Network UK, The National Cancer Research Institute (NCRI) UK. We are grateful for support of NIHR funding to the NIHR Biomedical Research Centre at The Institute of Cancer Research and The Royal Marsden NHS Foundation Trust.

The Prostate Cancer Program of Cancer Council Victoria also acknowledge grant support from The National Health and Medical Research Council, Australia (126402, 209057, 251533, 396414, 450104, 504700, 504702, 504715, 623204, 940394, 614296,), VicHealth, Cancer Council Victoria, The Prostate Cancer Foundation of Australia, The Whitten Foundation, PricewaterhouseCoopers, and Tattersall’s. EAO, DMK, and EMK acknowledge the Intramural Program of the National Human Genome Research Institute for their support.

Genotyping of the OncoArray was funded by the US National Institutes of Health (NIH) [U19 CA 148537 for ELucidating Loci Involved in Prostate cancer SuscEptibility (ELLIPSE) project and X01HG007492 to the Center for Inherited Disease Research (CIDR) under contract number HHSN268201200008I]. Additional analytic support was provided by NIH NCI U01 CA188392 (PI: Schumacher).

Funding for the iCOGS infrastructure came from: the European Community's Seventh Framework Programme under grant agreement n° 223175 (HEALTH-F2-2009-223175) (COGS), Cancer Research UK (C1287/A10118, C1287/A 10710, C12292/A11174, C1281/A12014, C5047/A8384, C5047/A15007, C5047/A10692, C8197/A16565), the National Institutes of Health (CA128978) and Post-Cancer GWAS initiative (1U19 CA148537, 1U19 CA148065 and 1U19 CA148112 - the GAME-ON initiative), the Department of Defence (W81XWH-10-1-0341), the Canadian Institutes of Health Research (CIHR) for the CIHR Team in Familial Risks of Breast Cancer, Komen Foundation for the Cure, the Breast Cancer Research Foundation, and the Ovarian Cancer Research Fund.

*BPC3*

The BPC3 was supported by the U.S. National Institutes of Health, National Cancer Institute (cooperative agreements U01-CA98233 to D.J.H., U01-CA98710 to S.M.G., U01-CA98216 to E.R., and U01-CA98758 to B.E.H., and Intramural Research Program of NIH/National Cancer Institute, Division of Cancer Epidemiology and Genetics).

*CAPS*

CAPS GWAS study was supported by the Cancer Risk Prediction Center (CRisP; www.crispcenter.org), a Linneus Centre (Contract ID 70867902) financed by the Swedish Research Council, (grant no K2010-70X-20430-04-3), the Swedish Cancer Foundation (grant no 09-0677), the Hedlund Foundation, the Soederberg Foundation, the Enqvist Foundation, ALF funds from the Stockholm County Council. Stiftelsen Johanna Hagstrand och Sigfrid Linner's Minne, Karlsson's Fund for urological and surgical research.

*PEGASUS*

PEGASUS was supported by the Intramural Research Program, Division of Cancer Epidemiology and Genetics, National Cancer Institute, National Institutes of Health.

Ovarian Cancer Association Consortium (OCAC)

We used published genome-wide association data reported by the Ovarian Cancer Association Consortium (OCAC). Summary genetic association data were obtained on 25,509 women with epithelial ovarian cancer and 40,941 controls of European descent. Genotyping was performed using the Illumina Custom Infinium array (OncoArray). The dataset comprises 63 project or case-control sets recruited from 14 countries representing participants of European ancestry. GWAS datasets were adjusted for population substructure and summary estimates were meta-analysed using an inverse-variance fixed-effects approach. Details on the genotyping protocol, GWAS and meta-analysis have been previously described[5,6].

Ethical approval from relevant research ethics committees was obtained for all studies in OCAC and written, informed consent was obtained from all participants in these studies.

International Lung Cancer Consortium (ILCCO)

We retrieved publicly available GWAS summary data on lung cancer from the International Lung Cancer Consortium (ILCCO) (11,348 lung cancer cases and 15,861 controls of European ancestry) which has been deposited on MRBase[7]. Details on the genotyping protocol, GWAS and meta-analysis have been previously described[7].

The Genetic and Epidemiology of Colorectal Cancer Consortium (GECCO), the Colorectal Cancer Transdisciplinary Study (CORECT), and the Colon Cancer Family Registry (CCFR) consortia (GECCO-CORECT-CCFR)

A recently published large GWAS of almost 126,000 participants of European ancestry from the GECCO, CORECT and CCFR consortia provided the genetic effects of the serum folate instruments on the risk of colorectal cancer (58,221 cases and 67,694 controls). GWAS were adjusted for age, sex, study and principle components to adjust for population substructure. Further details relating to the genotyping protocols, GWAS and meta-analysis have been previously described[8]

**Authors funding and acknowledgements**

***Co-authors from the GECCO, CORECT and CCFR consortia:***

*GECCO*

Albert de la Chapelle1, Amit D. Joshi2,41, Andrea Burnett-Hartman3, Andrea Gsur4, Andrew T. Chan5,42,59,65,67,68, Antonia Trichopoulou6,43, Barbara L. Banbury7, Bethany Van Guelpen8, Carlo La Vecchia9, Catherine M. Tangen10, Christina Bamia6,43, Christopher I. Li7, Conghui Qu7, D Timothy Bishop11, Dallas R. English12,44, David J. Hunter2,45, Elizabeth A. Platz13, Ellen Kampman14, Emily White7,46, Franzel JB. van Duijnhoven14, Giovanna Masala15, Graham G. Giles16,47,60, Hansong Wang17, Heather Hampel18, Heiner Boeing19, Henk van Kranen20, Hermann Brenner21,48,61, Jenny Chang-Claude22,49, Jeroen R. Huyghe7, John D. Potter7, Kala Visvanathan13, Li Hsu7,50, Lori C. Sakoda23,51, Ludmila Vodickova24,52,62, Marc J. Gunter25, Maria-Dolores Chirlaque26,53, Martha L. Slattery27, Michael Hoffmeister21, Michael O. Woods28, Miguel Rodríguez-Barranco29,54, Mingyang Song30,55,63, N Charlotte Onland-Moret31, Pavel Vodicka24,52,62, Peter T. Campbell32, Phyllis J. Goodman10, Roger L. Milne16,47,60, Sabina Sieri33, Sébastien Küry34, Sergi Castellví-Bel35, Sjoerd G. Elias31, Sonja I. Berndt36, Stephen J. Chanock36, Temitope O. Keku37, Tilman Kuhn22, Timothy J. Key38, Ulrike Peters7,56, Vicente Martín26,57, Victor Moreno39,54,64,66, Vittorio Perduca40,58

1. Department of Cancer Biology and Genetics and the Comprehensive Cancer Center, The Ohio State University, Columbus, Ohio, USA.
2. Department of Epidemiology, Harvard T.H. Chan School of Public Health, Harvard University, Boston, Massachusetts, USA.
3. Institute for Health Research, Kaiser Permanente Colorado, Denver, Colorado, USA.
4. Institute of Cancer Research, Department of Medicine I, Medical University Vienna, Vienna, Austria.
5. Division of Gastroenterology, Massachusetts General Hospital and Harvard Medical School, Boston, Massachusetts, USA.
6. Hellenic Health Foundation, Athens, Greece.
7. Public Health Sciences Division, Fred Hutchinson Cancer Research Center, Seattle, Washington, USA.
8. Department of Radiation Sciences, Oncology Unit, Umeå University, Umeå, Sweden.
9. Hellenic Health Foundation,Athens, Greece; Department of Clinical Sciences and Community Health, Università degli Studi di Milano, Milan, Italy.
10. SWOG Statistical Center, Fred Hutchinson Cancer Research Center, Seattle, Washington, USA.
11. Leeds Institute of Cancer and Pathology, University of Leeds, Leeds, UK.
12. Centre for Epidemiology and Biostatistics, Melbourne School of Population and Global Health, The University of Melbourne, Melbourne, Victoria, Australia.
13. Department of Epidemiology, Johns Hopkins Bloomberg School of Public Health, Baltimore, Maryland, USA.
14. Division of Human Nutrition, Wageningen University and Research, Wageningen, The Netherlands.
15. Cancer Risk Factors and Life-Style Epidemiology Unit, Institute of Cancer Research, Prevention and Clinical Network - ISPRO, Florence, Italy.
16. Cancer Epidemiology Division, Cancer Council Victoria, Melbourne, Victoria, Australia.
17. University of Hawaii Cancer Center, Honolulu, Hawaii, USA.
18. Division of Human Genetics, Department of Internal Medicine, The Ohio State University Comprehensive Cancer Center, Columbus, Ohio, USA.
19. Department of Epidemiology, German Institute of Human Nutrition (DIfE), Potsdam-Rehbrücke, Germany.
20. National Institute for Public Health and the Environment (RIVM), Bilthoven, The Netherlands.
21. Division of Clinical Epidemiology and Aging Research, German Cancer Research Center (DKFZ), Heidelberg, Germany.
22. Division of Cancer Epidemiology, German Cancer Research Center (DKFZ), Heidelberg, Germany.
23. Division of Research, Kaiser Permanente Northern California, Oakland, California, USA.
24. Department of Molecular Biology of Cancer, Institute of Experimental Medicine of the Czech Academy of Sciences, Prague, Czech Republic.
25. Nutrition and Metabolism Section, International Agency for Research on Cancer, World Health Organization, Lyon, France.
26. CIBER Epidemiología y Salud Pública (CIBERESP), Madrid, Spain.
27. Department of Internal Medicine, University of Utah, Salt Lake City, Utah, USA.
28. Memorial University of Newfoundland, Discipline of Genetics, St. John's, Canada.
29. Escuela Andaluza de Salud Pública, Instituto de Investigación Biosanitaria ibs.GRANADA, Hospitales Universitarios de Granada/Universidad de Granada, Granada, Spain.
30. Clinical and Translational Epidemiology Unit, Massachusetts General Hospital and Harvard Medical School, Boston, Massachusetts, USA.
31. Julius Center for Health Sciences and Primary Care, University Medical Center Utrecht, Utrecht, The Netherlands.
32. Behavioral and Epidemiology Research Group, American Cancer Society, Atlanta, Georgia, USA.
33. Epidemiology and Prevention Unit, Fondazione IRCCS Istituto Nazionale dei Tumori, Milan, Italy.
34. Service de Génétique Médicale, Centre Hospitalier Universitaire (CHU) Nantes, Nantes, France.
35. Gastroenterology Department, Hospital Clínic, Institut d'Investigacions Biomèdiques August Pi i Sunyer (IDIBAPS), Centro de Investigación Biomédica en Red de Enfermedades Hepáticas y Digestivas (CIBEREHD), University of Barcelona, Barcelona, Spain.
36. Division of Cancer Epidemiology and Genetics, National Cancer Institute, National Institutes of Health, Bethesda, Maryland, USA.
37. Center for Gastrointestinal Biology and Disease, University of North Carolina, Chapel Hill, North Carolina, USA.
38. Cancer Epidemiology Unit, Nuffield Department of Population Health, University of Oxford, Oxford, UK.
39. Oncology Data Analytics Program, Catalan Institute of Oncology-IDIBELL, L'Hospitalet de Llobregat, Barcelona, Spain.
40. Laboratoire de Mathématiques Appliquées MAP5 (UMR CNRS 8145), Université Paris Descartes, Paris, France.
41. Clinical and Translational Epidemiology Unit, Massachusetts General Hospital and Harvard Medical School, Boston, Massachusetts, USA.
42. Channing Division of Network Medicine, Brigham and Women's Hospital and Harvard Medical School, Boston, Massachusetts, USA.
43. WHO Collaborating Center for Nutrition and Health, Unit of Nutritional Epidemiology and Nutrition in Public Health, Deptartment of Hygiene, Epidemiology and Medical Statistics, School of Medicine, National and Kapodistrian University of Athens, Greece.
44. Cancer Epidemiology Division, Cancer Council Victoria, Melbourne, Victoria, Australia.
45. Nuffield Department of Population Health, University of Oxford, Oxford, UK.
46. Department of Epidemiology, University of Washington School of Public Health, Seattle, Washington, USA.
47. Centre for Epidemiology and Biostatistics, Melbourne School of Population and Global Health, The University of Melbourne, Melbourne, Victoria, Australia.
48. Division of Preventive Oncology, German Cancer Research Center (DKFZ) and National Center for Tumor Diseases (NCT), Heidelberg, Germany.
49. University Medical Centre Hamburg-Eppendorf, University Cancer Centre Hamburg (UCCH), Hamburg, Germany.
50. Department of Biostatistics, University of Washington, Seattle, Washington, USA
51. Public Health Sciences Division, Fred Hutchinson Cancer Research Center, Seattle, Washington, USA.
52. Institute of Biology and Medical Genetics, First Faculty of Medicine, Charles University, Prague, Czech Republic.
53. Department of Epidemiology, Regional Health Council, IMIB-Arrixaca, Murcia, Spain.
54. CIBER de Epidemiología y Salud Pública (CIBERESP), Madrid, Spain.
55. Division of Gastroenterology, Massachusetts General Hospital and Harvard Medical School, Boston, Massachusetts, USA.
56. Department of Epidemiology, University of Washington, Seattle, Washington, USA
57. Biomedicine Institute (IBIOMED), University of León, León, Spain.
58. CESP (Inserm U1018), Facultés de Medicine Université Paris-Sud, UVSQ, Université Paris-Saclay, Gustave Roussy, Villejuif, France.
59. Clinical and Translational Epidemiology Unit, Massachusetts General Hospital and Harvard Medical School, Boston, Massachusetts, USA.
60. Precision Medicine, School of Clinical Sciences at Monash Health, Monash University, Clayton, Victoria, Australia.
61. German Cancer Consortium (DKTK), German Cancer Research Center (DKFZ), Heidelberg, Germany.
62. Faculty of Medicine and Biomedical Center in Pilsen, Charles University, Pilsen, Czech Republic.
63. Department of Nutrition, Harvard T.H. Chan School of Public Health, Harvard University, Boston, Massachusetts, USA.
64. Department of Clinical Sciences, Faculty of Medicine, University of Barcelona, Barcelona, Spain.
65. Broad Institute of Harvard and MIT, Cambridge, Massachusetts, USA.
66. ONCOBEL Program, Bellvitge Biomedical Research Institute (IDIBELL), L'Hospitalet de Llobregat, Barcelona, Spain.
67. Department of Epidemiology, Harvard T.H. Chan School of Public Health, Harvard University, Boston, Massachusetts, USA.
68. Department of Immunology and Infectious Diseases, Harvard T.H. Chan School of Public Health, Harvard University, Boston, Massachusetts, USA.

*CORECT*

Alicja Wolk1, Anna H. Wu2, Annika Lindblom3,21, Clemens Schafmayer4, Cornelia M. Ulrich5, Demetrius Albanes6, Gad Rennert7,22,26, Hyeong Rok Kim8, Jane C. Figueiredo9,23, Jochen Hampe10, Juergen Boehm5, Kenneth Offit11,24, Li Li12, Min-Ho Shin13, Paul D. P. Pharoah14, Sang Hee Cho15, Stephanie J. Weinstein6, Stephanie L. Schmit16, Stephen B. Gruber17, Sun-Seog Kweon13,25, Susanna C. Larsson1, Volker Arndt18, Wei Zheng19, Zsofia K. Stadler20

1. Institute of Environmental Medicine, Karolinska Institutet, Stockholm, Sweden.
2. University of Southern California, Preventative Medicine, Los Angeles, California, USA.
3. Department of Clinical Genetics, Karolinska University Hospital, Stockholm, Sweden.
4. Department5 of General Surgery, University Hospital Rostock, Rostock, Germany.
5. Huntsman Cancer Institute and Department of Population Health Sciences, University of Utah, Salt Lake City, Utah, USA.
6. Division of Cancer Epidemiology and Genetics, National Cancer Institute, National Institutes of Health, Bethesda, Maryland, USA.
7. Department of Community Medicine and Epidemiology, Lady Davis Carmel Medical Center, Haifa, Israel.
8. Department of Surgery, Chonnam National University Hwasun Hospital and Medical School, Hwasun, Korea.
9. Department of Medicine, Samuel Oschin Comprehensive Cancer Institute, Cedars-Sinai Medical Center, Los Angeles, CA, USA.
10. Department of Medicine I, University Hospital Dresden, Technische Universität Dresden (TU Dresden), Dresden, Germany.
11. Clinical Genetics Service, Department of Medicine, Memorial Sloan-Kettering Cancer Center, New York, New York, USA.
12. Department of Family Medicine, University of Virginia, Charlottesville, Virginia, USA.
13. Department of Preventive Medicine, Chonnam National University Medical School, Gwangju, Korea.
14. Department of Public Health and Primary Care, University of Cambridge, Cambridge, UK.
15. Department of Hematology-Oncology, Chonnam National University Hospital, Hwasun, South Korea.
16. Department of Cancer Epidemiology, H. Lee Moffitt Cancer Center and Research Institute, Tampa, Florida, USA.
17. Department of Preventive Medicine & USC Norris Comprehensive Cancer Center, Keck School of Medicine, University of Southern California, Los Angeles, California, USA.
18. Division of Clinical Epidemiology and Aging Research, German Cancer Research Center (DKFZ), Heidelberg, Germany.
19. Division of Epidemiology, Department of Medicine, Vanderbilt-Ingram Cancer Center, Vanderbilt Epidemiology Center, Vanderbilt University School of Medicine, Nashville, Tennessee, USA.
20. Department of Medicine, Memorial Sloan Kettering Cancer Center, New York, New York, USA.
21. Department of Molecular Medicine and Surgery, Karolinska Institutet, Stockholm, Sweden.
22. Ruth and Bruce Rappaport Faculty of Medicine, Technion-Israel Institute of Technology, Haifa, Israel.
23. Department of Preventive Medicine, Keck School of Medicine, University of Southern California, Los Angeles, California, USA.
24. Department of Medicine, Weill Cornell Medical College, New York, New York, USA.
25. Jeonnam Regional Cancer Center, Chonnam National University Hwasun Hospital, Hwasun, Korea.
26. Clalit National Cancer Control Center, Haifa, Israel.

*CCFR*

Daniel D. Buchanan1,10,13, Graham Casey2, Jane C. Figueiredo3,11, Loic Le Marchand4, Mark A. Jenkins5, Noralane M. Lindor6, Polly A. Newcomb7,12, Stephen N. Thibodeau8, Steven J. Gallinger9

1. Colorectal Oncogenomics Group, Department of Clinical Pathology, The University of Melbourne, Parkville, Victoria 3010 Australia
2. Center for Public Health Genomics, University of Virginia, Charlottesville, Virginia, USA.
3. Department of Medicine, Samuel Oschin Comprehensive Cancer Institute, Cedars-Sinai Medical Center, Los Angeles, CA, USA.
4. University of Hawaii Cancer Center, Honolulu, Hawaii, USA.
5. Centre for Epidemiology and Biostatistics, Melbourne School of Population and Global Health, The University of Melbourne, Melbourne, Victoria, Australia.
6. Department of Health Science Research, Mayo Clinic, Scottsdale, Arizona, USA.
7. Public Health Sciences Division, Fred Hutchinson Cancer Research Center, Seattle, Washington, USA.
8. Division of Laboratory Genetics, Department of Laboratory Medicine and Pathology, Mayo Clinic, Rochester, Minnesota, USA.
9. Lunenfeld Tanenbaum Research Institute, Mount Sinai Hospital, University of Toronto, Toronto, Ontario, Canada.
10. University of Melbourne Centre for Cancer Research, Victorian Comprehensive Cancer Centre, Parkville, Victoria 3010 Australia
11. Department of Preventive Medicine, Keck School of Medicine, University of Southern California, Los Angeles, California, USA.
12. School of Public Health, University of Washington, Seattle, Washington, USA.
13. Genetic Medicine and Family Cancer Clinic, The Royal Melbourne Hospital, Parkville, Victoria, Australia.

Funding

Genetics and Epidemiology of Colorectal Cancer Consortium (GECCO): National Cancer Institute, National Institutes of Health, U.S. Department of Health and Human Services (U01 CA164930, U01 CA137088, R01 CA059045, U01 CA164930, R21 CA191312).

ASTERISK: a Hospital Clinical Research Program (PHRC-BRD09/C) from the University Hospital Center of Nantes (CHU de Nantes) and supported by the Regional Council of Pays de la Loire, the Groupement des Entreprises Françaises dans la Lutte contre le Cancer (GEFLUC), the Association Anne de Bretagne Génétique and the Ligue Régionale Contre le Cancer (LRCC).

The ATBC Study is supported by the Intramural Research Program of the U.S. National Cancer Institute, National Institutes of Health, and by U.S. Public Health Service contract HHSN261201500005C from the National Cancer Institute, Department of Health and Human Services.

CLUE II: This research was funded by the American Institute for Cancer Research and the Maryland Cigarette Restitution Fund at Johns Hopkins, and the NCI (P30 CA006973 to W.G. Nelson).

COLO2&3: National Institutes of Health (R01 CA60987).

ColoCare: This work was supported by the National Institutes of Health (grant numbers R01 CA189184 (Li/Ulrich), U01 CA206110 (Ulrich/Li/Siegel/Figueireido/Colditz, 2P30CA015704- 40 (Gilliland), R01 CA207371 (Ulrich/Li)), the Matthias Lackas-Foundation, the German Consortium for Translational Cancer Research, and the EU TRANSCAN initiative.

The Colon Cancer Family Registry (CCFR, www.coloncfr.org) was supported in part by funding from the National Cancer Institute (NCI), National Institutes of Health (NIH) (award U01 CA167551) and through U01/U24 cooperative agreements from NCI with the following CCFR centers: Australasian (CA074778 and CA097735), , Ontario (OFCCR) (CA074783), Seattle (SFCCR) (CA074794 (and R01 CA076366 to PAN)), USC Consortium (CA074799), Mayo Clinic (CA074800), and Hawaii (CA074806). Support for case ascertainment was provided in part from the Surveillance, Epidemiology, and End Results (SEER) Program and the following U.S. state cancer registries: AZ, CO, MN, NC, NH; and by the Victoria Cancer Registry (Australia) and Ontario Cancer Registry (Canada). Additional funding for the OFCCR/ARCTIC was through award GL201-043 from the Ontario Research Fund (to BWZ), award 112746 from the Canadian Institutes of Health Research (to TJH), through a Cancer Risk Evaluation (CaRE) Program grant from the Canadian Cancer Society (to SG), and through generous support from the Ontario Ministry of Research and Innovation. The SCCFR Illumina HumanCytoSNP array was supported through NCI award R01 CA076366 (to PAN). The CCFR Set-1 (Illumina 1M/1M-Duo) and Set-2 (Illumina Omni1-Quad) scans were supported by NIH awards U01 CA122839 and R01 CA143247 (to GC). The CCFR Set-3 (Affymetrix Axiom CORECT Set array) was supported by NIH award U19 CA148107 and R01 CA81488 (to SBG). The CCFR Set-4 (Illumina OncoArray 600K SNP array) was supported by NIH award U19 CA148107 (to SBG) and by the Center for Inherited Disease Research (CIDR), which is funded by the NIH to the Johns Hopkins University, contract number HHSN268201200008I. Colon Cancer Family Registry (CCFR): The content of this manuscript does not necessarily reflect the views or policies of the NIH or any of the collaborating centers in the CCFR, nor does mention of trade names, commercial products, or organizations imply endorsement by the US Government, any cancer registry, or the CCFR.

COLON: The COLON study is sponsored by Wereld Kanker Onderzoek Fonds, including funds from grant 2014/1179 as part of the World Cancer Research Fund International Regular Grant Programme, by Alpe d’Huzes and the Dutch Cancer Society (UM 2012–5653, UW 2013-5927, UW2015-7946), and by TRANSCAN (JTC2012-MetaboCCC, JTC2013-FOCUS). The Nqplus study is sponsored by a ZonMW investment grant (98-10030); by PREVIEW, the project PREVention of diabetes through lifestyle intervention and population studies in Europe and around the World (PREVIEW) project which received funding from the European Union Seventh Framework Programme (FP7/2007–2013) under grant no. 312057; by funds from TI Food and Nutrition (cardiovascular health theme), a public–private partnership on precompetitive research in food and nutrition; and by FOODBALL, the Food Biomarker Alliance, a project from JPI Healthy Diet for a Healthy Life.

Colorectal Cancer Transdisciplinary (CORECT) Study: The CORECT Study was supported by the National Cancer Institute, National Institutes of Health (NCI/NIH), U.S. Department of Health and Human Services (grant numbers U19 CA148107, R01 CA81488, P30 CA014089, R01 CA197350,; P01 CA196569; R01 CA201407) and National Institutes of Environmental Health Sciences, National Institutes of Health (grant number T32 ES013678).

CORSA: “Österreichische Nationalbank Jubiläumsfondsprojekt” (12511) and Austrian Research Funding Agency (FFG) grant 829675.

CPS-II: The American Cancer Society funds the creation, maintenance, and updating of the Cancer Prevention Study-II (CPS-II) cohort. This study was conducted with Institutional Review Board approval.

CRCGEN: Colorectal Cancer Genetics & Genomics, Spanish study was supported by Instituto de Salud Carlos III, co-funded by FEDER funds –a way to build Europe– (grants PI14-613 and PI09-1286), Agency for Management of University and Research Grants (AGAUR) of the Catalan Government (grant 2017SGR723), and Junta de Castilla y León (grant LE22A10-2). Sample collection of this work was supported by the Xarxa de Bancs de Tumors de Catalunya sponsored by Pla Director d’Oncología de Catalunya (XBTC), Plataforma Biobancos PT13/0010/0013 and ICOBIOBANC, sponsored by the Catalan Institute of Oncology.

Czech Republic CCS: This work was supported by the Grant Agency of the Czech Republic (grants CZ GA CR: GAP304/10/1286 and 1585) and by the Grant Agency of the Ministry of Health of the Czech Republic (grants AZV 15-27580A and AZV 17-30920A).

DACHS: This work was supported by the German Research Council (BR 1704/6-1, BR 1704/6-3, BR 1704/6-4, CH 117/1-1, HO 5117/2-1, HE 5998/2-1, KL 2354/3-1, RO 2270/8-1 and BR 1704/17-1), the Interdisciplinary Research Program of the National Center for Tumor Diseases (NCT), Germany, and the German Federal Ministry of Education and Research (01KH0404, 01ER0814, 01ER0815, 01ER1505A and 01ER1505B).

DALS: National Institutes of Health (R01 CA48998 to M. L. Slattery).

EDRN: This work is funded and supported by the NCI, EDRN Grant (U01 CA 84968-06).

EPIC: The coordination of EPIC is financially supported by the European Commission (DGSANCO) and the International Agency for Research on Cancer. The national cohorts are supported by Danish Cancer Society (Denmark); Ligue Contre le Cancer, Institut Gustave Roussy, Mutuelle Générale de l’Education Nationale, Institut National de la Santé et de la Recherche Médicale (INSERM) (France); German Cancer Aid, German Cancer Research Center (DKFZ), Federal Ministry of Education and Research (BMBF), Deutsche Krebshilfe, Deutsches Krebsforschungszentrum and Federal Ministry of Education and Research (Germany); the Hellenic Health Foundation (Greece); Associazione Italiana per la Ricerca sul Cancro-AIRCItaly and National Research Council (Italy); Dutch Ministry of Public Health, Welfare and Sports (VWS), Netherlands Cancer Registry (NKR), LK Research Funds, Dutch Prevention Funds, Dutch ZON (Zorg Onderzoek Nederland), World Cancer Research Fund (WCRF), Statistics Netherlands (The Netherlands); ERC-2009-AdG 232997 and Nordforsk, Nordic Centre of Excellence programme on Food, Nutrition and Health (Norway); Health Research Fund (FIS), PI13/00061 to Granada, PI13/01162 to EPIC-Murcia, Regional Governments of Andalucía, Asturias, Basque Country, Murcia and Navarra, ISCIII RETIC (RD06/0020) (Spain); Swedish Cancer Society, Swedish Research Council and County Councils of Skåne and Västerbotten (Sweden); Cancer Research UK (14136 to EPIC-Norfolk; C570/A16491 and C8221/A19170 to EPIC-Oxford), Medical Research Council (1000143 to EPIC-Norfolk, MR/M012190/1 to EPICOxford) (United Kingdom).

EPICOLON: This work was supported by grants from Fondo de Investigación Sanitaria/FEDER (PI08/0024, PI08/1276, PS09/02368, P111/00219, PI11/00681, PI14/00173, PI14/00230, PI17/00509, 17/00878, Acción Transversal de Cáncer), Xunta de Galicia (PGIDIT07PXIB9101209PR), Ministerio de Economia y Competitividad (SAF07-64873, SAF 2010-19273, SAF2014-54453R), Fundación Científica de la Asociación Española contra el Cáncer (GCB13131592CAST), Beca Grupo de Trabajo “Oncología” AEG (Asociación Española de Gastroenterología), Fundación Privada Olga Torres, FP7 CHIBCHA Consortium, Agència de Gestió d’Ajuts Universitaris i de Recerca (AGAUR, Generalitat de Catalunya, 2014SGR135, 2014SGR255, 2017SGR21, 2017SGR653), Catalan Tumour Bank Network (Pla Director d’Oncologia, Generalitat de Catalunya), PERIS (SLT002/16/00398, Generalitat de Catalunya), CERCA Programme (Generalitat de Catalunya) and COST Action BM1206 and CA17118. CIBERehd is funded by the Instituto de Salud Carlos III.

ESTHER/VERDI. This work was supported by grants from the Baden-Württemberg Ministry of Science, Research and Arts and the German Cancer Aid.

Harvard cohorts (HPFS, NHS, PHS): HPFS is supported by the National Institutes of Health (P01 CA055075, UM1 CA167552, U01 CA167552, R01 CA137178, R01 CA151993, R35 CA197735, K07 CA190673, and P50 CA127003), NHS by the National Institutes of Health (R01 CA137178, P01 CA087969, UM1 CA186107, R01 CA151993, R35 CA197735, K07CA190673, and P50 CA127003) and PHS by the National Institutes of Health (R01 CA042182).

Hawaii Adenoma Study: NCI grants R01 CA72520.

HCES-CRC: the Hwasun Cancer Epidemiology Study–Colon and Rectum Cancer (HCES-CRC; grants from Chonnam National University Hwasun Hospital, HCRI15011-1).

Kentucky: This work was supported by the following grant support: Clinical Investigator Award from Damon Runyon Cancer Research Foundation (CI-8); NCI R01CA136726.

LCCS: The Leeds Colorectal Cancer Study was funded by the Food Standards Agency and Cancer Research UK Programme Award (C588/A19167).

MCCS cohort recruitment was funded by VicHealth and Cancer Council Victoria. The MCCS was further supported by Australian NHMRC grants 509348, 209057, 251553 and 504711 and by infrastructure provided by Cancer Council Victoria. Cases and their vital status were ascertained through the Victorian Cancer Registry (VCR) and the Australian Institute of Health and Welfare (AIHW), including the National Death Index and the Australian Cancer Database.

MEC: National Institutes of Health (R37 CA54281, P01 CA033619, and R01 CA063464).

MECC: This work was supported by the National Institutes of Health, U.S. Department of Health and Human Services (R01 CA81488 to SBG and GR).

MSKCC: The work at Sloan Kettering in New York was supported by the Robert and Kate Niehaus Center for Inherited Cancer Genomics and the Romeo Milio Foundation. Moffitt: This work was supported by funding from the National Institutes of Health (grant numbers R01 CA189184, P30 CA076292), Florida Department of Health Bankhead-Coley Grant 09BN-13, and the University of South Florida Oehler Foundation. Moffitt contributions were supported in part by the Total Cancer Care Initiative, Collaborative Data Services Core, and Tissue Core at the H. Lee Moffitt Cancer Center & Research Institute, a National Cancer Institute-designated Comprehensive Cancer Center (grant number P30 CA076292).

NCCCS I & II: We acknowledge funding support for this project from the National Institutes of Health, R01 CA66635 and P30 DK034987.

NFCCR: This work was supported by an Interdisciplinary Health Research Team award from the Canadian Institutes of Health Research (CRT 43821); the National Institutes of Health, U.S. Department of Health and Human Serivces (U01 CA74783); and National Cancer Institute of Canada grants (18223 and 18226). The authors wish to acknowledge the contribution of Alexandre Belisle and the genotyping team of the McGill University and Génome Québec Innovation Centre, Montréal, Canada, for genotyping the Sequenom panel in the NFCCR samples. Funding was provided to Michael O. Woods by the Canadian Cancer Society Research Institute.

NSHDS: Swedish Cancer Society; Cancer Research Foundation in Northern Sweden; Swedish Research Council; J C Kempe Memorial Fund; Faculty of Medicine, Umeå University, Umeå, Sweden; and Cutting-Edge Research Grant from the County Council of Västerbotten, Sweden.

OFCCR: The Ontario Familial Colorectal Cancer Registry was supported in part by the National Cancer Institute (NCI) of the National Institutes of Health (NIH) under award U01 CA167551 and award U01/U24 CA074783 (to SG). Additional funding for the OFCCR and ARCTIC testing and genetic analysis was through and a Canadian Cancer Society CaRE (Cancer Risk Evaluation) program grant and Ontario Research Fund award GL201-043 (to BWZ), through the Canadian Institutes of Health Research award 112746 (to TJH), and through generous support from the Ontario Ministry of Research and Innovation.

OSUMC: OCCPI funding was provided by Pelotonia and HNPCC funding was provided by the NCI (CA16058 and CA67941).

PLCO: Intramural Research Program of the Division of Cancer Epidemiology and Genetics and supported by contracts from the Division of Cancer Prevention, National Cancer Institute, NIH, DHHS. Funding was provided by National Institutes of Health (NIH), Genes, Environment and Health Initiative (GEI) Z01 CP 010200, NIH U01 HG004446, and NIH GEI U01 HG 004438.

REACH: This work was supported by the National Cancer Institute (grant P01 CA074184 to J.D.P. and P.A.N., grants R01 CA097325, R03 CA153323, and K05 CA152715 to P.A.N., and the National Center for Advancing Translational Sciences at the National Institutes of Health (grant KL2 TR000421 to A.N.B.-H.)

SCCFR: The Seattle Colon Cancer Family Registry was supported in part by the National Cancer Institute (NCI) of the National Institutes of Health (NIH) under awards U01 CA167551. Additional support for the SFCCR, Postmenopausal Hormones and Colon Cancer (PMH) study and the SCCFR Illumina HumanCytoSNP array were through NCI/NIH awards U01/U24 CA074794 and R01 CA076366 (to PAN).

SEARCH: The University of Cambridge has received salary support in respect of PDPP from the NHS in the East of England through the Clinical Academic Reserve. Cancer Research UK (C490/A16561); the UK National Institute for Health Research Biomedical Research Centres at the University of Cambridge.

SELECT: Research reported in this publication was supported in part by the National Cancer Institute of the National Institutes of Health under Award Numbers U10 CA37429 (CD Blanke), and UM1 CA182883 (CM Tangen/IM Thompson). The content is solely the responsibility of the authors and does not necessarily represent the official views of the National Institutes of Health.

SMS: This work was supported by the National Cancer Institute (grant P01 CA074184 to J.D.P. and P.A.N., grants R01 CA097325, R03 CA153323, and K05 CA152715 to P.A.N., and the National Center for Advancing Translational Sciences at the National Institutes of Health (grant KL2 TR000421 to A.N.B.-H.)

The Swedish Low-risk Colorectal Cancer Study: The study was supported by grants from the Swedish research council; K2015-55X-22674-01-4, K2008-55X-20157-03-3, K2006-72X-20157-01-2 and the Stockholm County Council (ALF project).

Swedish Mammography Cohort and Cohort of Swedish Men: This work is supported by the Swedish Research Council /Infrastructure grant, the Swedish Cancer Foundation, and the Karolinska Institute´s Distinguished Professor Award to Alicja Wolk.

UK Biobank: This research has been conducted using the UK Biobank Resource under Application Number 8614

VITAL: National Institutes of Health (K05 CA154337).

Women’s Health Initiative: The WHI program is funded by the National Heart, Lung, and Blood Institute, National Institutes of Health, U.S. Department of Health and Human Services through contracts HHSN268201100046C, HHSN268201100001C, HHSN268201100002C, HHSN268201100003C, HHSN268201100004C, and HHSN271201100004C.

Acknowledgements:

ASTERISK: We are very grateful to Dr. Bruno Buecher without whom this project would not have existed. We also thank all those who agreed to participate in this study, including the patients and the healthy control persons, as well as all the physicians, technicians and students.

CLUE II: We appreciate the continued efforts of the staff members at the Johns Hopkins George W. Comstock Center for Public Health Research and Prevention in the conduct of the CLUE II study. Cancer incidence data for CLUE were provided by the Maryland Cancer Registry, Center for Cancer Surveillance and Control, Maryland Department of Health, 201 W. Preston Street, Room 400, Baltimore, MD 21201, http://phpa.dhmh.maryland.gov/cancer, 410-767-4055. We acknowledge the State of Maryland, the Maryland Cigarette Restitution Fund, and the National Program of Cancer Registries of the Centers for Disease Control and Prevention for the funds that support the collection and availability of the cancer registry data.

COLON and NQplus: the authors would like to thank the COLON and NQplus investigators at Wageningen University & Research and the involved clinicians in the participating hospitals.

CCFR: The Colon CFR graciously thanks the generous contributions of their 42,505 study participants, dedication of study staff, and the financial support from the U.S. National Cancer Institute, without which this important registry would not exist.

CORSA: We kindly thank all those who contributed to the screening project Burgenland against CRC. Furthermore, we are grateful to Doris Mejri and Monika Hunjadi for laboratory assistance.

CPS-II: The authors thank the CPS-II participants and Study Management Group for their invaluable contributions to this research. The authors would also like to acknowledge the contribution to this study from central cancer registries supported through the Centers for Disease Control and Prevention National Program of Cancer Registries, and cancer registries supported by the National Cancer Institute Surveillance Epidemiology and End Results program.

Czech Republic CCS: We are thankful to all clinicians in major hospitals in the Czech Republic, without whom the study would not be practicable. We are also sincerely grateful to all patients participating in this study.

DACHS: We thank all participants and cooperating clinicians, and Ute Handte-Daub, Utz Benscheid, Muhabbet Celik and Ursula Eilber for excellent technical assistance.

EDRN: We acknowledge all the following contributors to the development of the resource: University of Pittsburgh School of Medicine, Department of Gastroenterology, Hepatology and Nutrition: Lynda Dzubinski; University of Pittsburgh School of Medicine, Department of Pathology: Michelle Bisceglia; and University of Pittsburgh School of Medicine, Department of Biomedical Informatics.

EPICOLON: We are sincerely grateful to all patients participating in this study who were recruited as part of the EPICOLON project. We acknowledge the Spanish National DNA Bank, the Barcelona CRC screening program, the Biobank core facility of Hospital Clínic–IDIBAPS and the Biobanco Vasco para la Investigación/O+ehun-Hospital Donostia for the availability of the samples. The work was carried out (in part) at the Esther Koplowitz Centre, Barcelona.

Harvard cohorts (HPFS, NHS, PHS): The study protocol was approved by the institutional review boards of the Brigham and Women’s Hospital and Harvard T.H. Chan School of Public Health, and those of participating registries as required. We would like to thank the participants and staff of the HPFS, NHS and PHS for their valuable contributions as well as the following state cancer registries for their help: AL, AZ, AR, CA, CO, CT, DE, FL, GA, ID, IL, IN, IA, KY, LA, ME, MD, MA, MI, NE, NH, NJ, NY, NC, ND, OH, OK, OR, PA, RI, SC, TN, TX, VA, WA, WY. The authors assume full responsibility for analyses and interpretation of these data.

Kentucky: We would like to acknowledge the staff at the Kentucky Cancer Registry.

LCCS: We acknowledge the contributions of Jennifer Barrett, Robin Waxman, Gillian Smith and Emma Northwood in conducting this study.

NCCCS I & II: We would like to thank the study participants, and the NC Colorectal Cancer Study staff.

NSHDS: NSHDS investigators thank the cohort participants, the Biobank Research Unit at Umeå University and Biobanken Norr at Region Västerbotten. The research was supported by Biobank Sweden and through funding from the Swedish Research Council; the Swedish Cancer Society; Region Västerbotten; the Lion’s Cancer Research Foundation, the Faculty of Medicine, and Insamlingsstiftelsen, all at Umeå University; and the Margareta Dannborg Memorial Fund.

PLCO: The authors thank the PLCO Cancer Screening Trial screening center investigators and the staff from Information Management Services Inc and Westat Inc. Most importantly, we thank the study participants for their contributions that made this study possible.

The SCCFR and the PMH study graciously thanks the generous contributions of their study participants, dedication of study staff, and the financial support from the U.S. National Cancer Institute, without which this important research was not possible. The content of this manuscript does not necessarily reflect the views or policies of the NIH or any of the collaborating centers in the CCFR, nor does mention of trade names, commercial products, or organizations imply endorsement by the US Government, any cancer registry, or the CCFR.

SEARCH: We thank the SEARCH team.

SELECT: We thank the research and clinical staff at the sites that participated on SELECT study, without whom the trial would not have been successful. We are also grateful to the 35,533 dedicated men who participated in SELECT.

Women’s Health Initiative: The authors thank the WHI investigators and staff for their dedication, and the study participants for making the program possible. A full listing of WHI investigators can be found at: http://www.whi.org/researchers/Documents%20%20Write%20a%20Paper/WHI%20Investigator%20Short%20List.pdf

### Statistical Information

Conversion of instrument GWAS effect estimates to the SD scale

The sample mean and SD on the original scale (serum folate nmol/L) was extracted from Shane *et al.*[9] for males and females separately. The mean and SD were pooled to derive an overall mean and SD for all participants included in the GWAS using the following equations[10]:

$$Combined Sample Mean= \frac{N_{1}M_{1}+N_{2}M_{2}}{N_{1}+N_{2}}$$

$$Combined Sample SD= \sqrt{\frac{\left( N_{1}-1 \right){SD}_{1}^{2}+\left( N_{2}-1 \right){SD}_{2}^{2}+\frac{N_{1}N_{2}}{N_{1}+N_{2}}\left( M_{1}^{2}+M_{2}^{2}-2M_{1}M_{2} \right)}{N_{1}+N_{2}-1}}$$

Where N_i_ is the sample size for study *i*, M_i_ is the sample mean for study *i* and SD_i_ is the sample SD of study *i*.

The sample SD was then converted to the Log_10_ scale using the following equations:

$$variance=log10\left( \frac{\left( {SD}^{2}+ {Mean}^{2} \right)}{{SD}^{2}} \right)$$

$$SD= \sqrt{variance}$$

We then divided the GWAS beta-coefficients and SE by the above SD, which is now on the logarithmic scale resulting in the effect estimates representing the change in SD of serum folate per each additional effect allele.

Strength of MR instruments

R^2^ representing the amount of variance in serum folate explained by each of the three SNPs used to construct our instruments were derived using the following:

$$R^{2}=\frac{2\times\beta^{2}\times MAF\times\left( 1-MAF \right)}{2 \times\beta^{2}\times MAF\times\left( 1-MAF \right)+\left( SE\left( \beta\right) \right)^{2} \times\left( 2 \times N \right)\times MAF\times\left( 1-MAF \right)}$$

Where β is the effect size (beta coefficient) for a given SNP, MAF is the minor allele frequency, SE(β) is the standard error of the effect size, and N is the sample size of the GWAS for the SNP-risk factor association.

$$FStatistic=R^2\times(N-1-k))/((1-R^2 ) \times k)$$

Where R^2^ is the proportion of variance explained in the risk factor by the genetic instrument, N is the sample size of the GWAS, *k* is the number of SNPs included in the instrument.
